# Model-based assessment of public health impact and cost-effectiveness of dengue vaccination following screening for prior exposure

**DOI:** 10.1101/367060

**Authors:** Guido España, Yutong Yao, Kathryn B. Anderson, Meagan C. Fitzpatrick, David L. Smith, Amy C. Morrison, Annelies Wilder-Smith, Thomas W. Scott, T. Alex Perkins

## Abstract

The tetravalent dengue vaccine CYD-TDV (Dengvaxia^®^) is the first licensed vaccine against dengue, but recent findings indicate an elevated risk of severe disease among vaccinees without prior dengue virus (DENV) exposure. The World Health Organization currently recommends CYD-TDV only for individuals with serological confirmation of past DENV exposure. Our objective was to evaluate the potential health impact and cost-effectiveness of vaccination following serological screening. To do so, we used an agent-based model to simulate DENV transmission with and without vaccination over a 10-year timeframe. Across a range of values for the proportion of vaccinees with prior DENV exposure, we projected the proportion of symptomatic and hospitalized cases averted as a function of the sensitivity and specificity of serological screening. Scenarios about the cost-effectiveness of screening and vaccination were chosen to be representative of Brazil and the Philippines. We found that public health impact depended primarily on sensitivity in high-transmission settings and on specificity in low-transmission settings. Cost-effectiveness could be achievable from the perspective of a public payer provided that sensitivity and the value of a disability-adjusted life-year were both high, but only in high-transmission settings. Requirements for reducing relative risk and achieving cost-effectiveness from an individual perspective were more restricted, due to the fact that those who test negative pay for screening but receive no benefit. Our results predict that cost-effectiveness could be achieved only in high-transmission areas of dengue-endemic countries with a relatively high per capita GDP, such as Panamá (13,680 USD), Brazil (8,649 USD), México (8,201 USD), or Thailand (5,807 USD). In conclusion, vaccination with CYD-TDV following serological screening could have a positive impact in some high-transmission settings, provided that screening is highly specific (to minimize individual harm), at least moderately sensitive (to maximize population benefit), and sufficiently inexpensive (depending on the setting).

**AUTHOR SUMMARY:** Among several viral diseases transmitted by *Aedes aegypti* mosquitoes, dengue imposes the greatest and most persistent burden on global health. Efforts to curb its spread would benefit greatly from the availability of an effective vaccine. Currently, the only licensed dengue vaccine, known as CYD-TDV or by the brand name Dengvaxia^®^, is only recommended for use in people who are known to have been exposed to dengue virus in the past. Because symptoms of dengue can range from severe to mild to imperceptible, using clinical history alone to assess whether a person was previously exposed is unreliable. Instead, serological assays, which measure a person’s immune response to dengue virus, are necessary to confirm whether a person was previously exposed. Because serological assays can be subject to substantial error, we used a simulation model to assess how impactful CYD-TDV vaccination would be under different scenarios about the accuracy of a serological assay and the intensity of transmission in a given area. We found that the health impact and cost-effectiveness of CYD-TDV vaccination depended on the accuracy of the serological assay, its cost, and the setting in which it is deployed.

## INTRODUCTION

A safe and effective dengue vaccine could have a major public health impact, as dengue causes approximately 9,000 deaths and between 50-100 million clinically apparent cases worldwide every year [1,2] and has a growing geographic distribution [3]. The first licensed dengue vaccine, CYD-TDV (Dengvaxia^®^), is a tetravalent, live-attenuated vaccine that was licensed in multiple countries after demonstrating efficacy against symptomatic disease in phase-III trials [4,5]. Protection has been hypothesized to derive primarily from the vaccine functioning as a “silent infection” [6]. Following their first natural infection subsequent to vaccination, this mechanism would result in vaccinees with prior dengue virus (DENV) exposure bypassing the elevated risk of severe disease typically associated with secondary infections. Modeling analyses [6,7] indicated that vaccination of nine-year-old children with CYD-TDV could be cost-effective in populations in which the majority of vaccinees have prior DENV exposure.

The downside of this mode of protection is an elevated risk of severe disease in vaccinees with no prior DENV exposure at the time of their first natural DENV infection [8]. Recent findings [9] confirmed this hypothesis, leading to an abrupt end to CYD-TDV use in the Philippines [10] and a revision of the World Health Organization’s (WHO) Strategic Advisory Group of Experts on Immunization recommendations in April 2018 on the use of the vaccine [11]. Vaccination with CYD-TDV is now recommended only for individuals with known prior DENV exposure [12–14]. Because DENV infection often results in asymptomatic infection or presents with mild, non-specific symptoms [15], an individual’s clinical history is a poor indicator of prior exposure. Thus, serological screening must play a role in any path forward for CYD-TDV or any other future dengue vaccines with similar characteristics. Reliable inference of prior DENV exposure based on serological data can be extremely challenging, however, due to cross-reactivity among DENV serotypes and among DENV and other flaviviruses [16,17].

To avoid elevating the risk of severe dengue by vaccinating a DENV-naïve individual, serological screening used to inform vaccination must have high specificity (i.e., probability that a DENV-naïve individual tests seronegative). At the same time, high sensitivity (i.e., probability that an individual with prior DENV exposure tests seropositive) is important for ensuring that people who could benefit from the vaccine will receive it. The balance of benefits and harms caused by vaccination with CYD-TDV following serological screening with a given sensitivity and specificity must also be weighed against the economic benefits and costs of such a strategy. Although a strategy of CYD-TDV vaccination following serological screening has been examined with mathematical modeling before [18,19], those analyses were restricted to a scenario in which the screening assay had perfect sensitivity and specificity. In practice, imperfect sensitivity and specificity [20], tradeoffs between sensitivity and specificity [21], and cost [22] all merit consideration in analyses of serological screening in CYD-TDV vaccination programs.

We applied an agent-based model of DENV transmission to identify the conditions under which a strategy of vaccination with CYD-TDV following serological screening (hereafter, referred to together as “the intervention”) would have positive impacts on health and be cost-effective. As with a previous study [7] involving this model and seven others, we focused our analysis on a strategy of routine intervention applied to a single age of nine years old. From both an individual and population perspective, we identified minimum requirements to achieve positive health impact and cost-effectiveness as a function of sensitivity, specificity, cost of serological screening, cost of vaccination, and prior DENV exposure among nine-year-olds (PE9). We focused on cost scenarios representative of Brazil and the Philippines, which have both licensed CYD-TDV but differ in terms of economic conditions.

## METHODS

### Model description

Our agent-based model of DENV transmission was previously described elsewhere [23]. This model has been previously used as part of a consortium of eight modeling groups to make projections of CYD-TDV impact in the absence of serological screening [7]. Despite differences with the other models, our model showed general agreement on projections of vaccination impact. In our model, humans and mosquitoes are represented by individual agents who interact with each other through mosquito blood-feeding at the household scale. The model assumes that transmission of any of the four DENV serotypes can occur whenever an infected mosquito blood-feeds on a susceptible human or a susceptible mosquito blood-feeds on an infected human. Infected humans acquire life-long immunity to the infecting serotype and temporary immunity to other serotypes to which they have not been previously exposed. Several model features are parameterized based on extensive data collection from Iquitos, Peru, including fine-scale patterns of human mobility [24], the demographic composition of households [25], and the geographic arrangement of residential, commercial, and other buildings [26]. Other model features were less well known a priori: the rate at which DENV was seeded into the population, the probability of an infectious mosquito infecting a susceptible human during blood-feeding, and the emergence rate of adult female mosquitoes. For a given simulation, we parameterized these features of the model by selecting a combination of parameter values that achieved a target value of the proportion of nine-year-olds with prior DENV exposure after 40 years of simulation, or PE_9_, as described in Appendix S1.

### Vaccination following serological screening

The vaccine implemented in our simulations acted as a silent DENV infection in the recipient, as has been assumed in previous CYD-TDV modeling assessments [6,7]. Because the vaccine is assumed to act as a silent infection, vaccination results in an elevated risk of severe disease among DENV-naïve vaccinees experiencing their first natural DENV infection, because secondary infections are associated with the highest probabilities of symptomatic disease conditional on infection and hospitalization conditional on symptomatic disease. In addition, we assumed a period of temporary cross-immunity after vaccination that waned over time. The level of protection and the waning period varied for individuals with and without previous exposure to DENV. Death was assumed to occur among a small proportion (0.0078) of cases of symptomatic disease. Because estimates of the rates of these outcomes are highly variable across study settings [27], we calibrated our model such that its outputs matched the most recent estimates of vaccine protection from clinical trials [9]. We did so by simulating a virtual trial [28] similar to the trials across a range of values of ten model parameters (Table 2) using a sequential importance sampling approach [29] and generalized additive models from the ‘mgcv’ [30] library in R[31], as described in Appendix S2. The best-fit model showed agreement with estimates of vaccine efficacy against symptomatic disease and hazard ratios for hospitalization stratified by age and baseline serostatus (Appendix S2).

Consistent with recently revised WHO recommendations [12], we simulated serological screening immediately prior to vaccination with CYD-TDV. We focused on a strategy of routine vaccination in which a proportion of children underwent serological screening, and vaccination in the event of a positive result, on their ninth birthday. One consequence of this strategy was that intervention coverage (i.e., the proportion of children screened) represents an upper limit on the proportion of vaccine-eligible children. Assuming that all vaccine-eligible children were vaccinated, the vaccination coverage (i.e., positive serological screening result and subsequent vaccination) was related to intervention coverage by

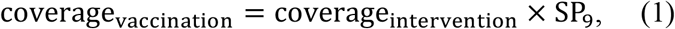

where SP9 is seropositivity among nine-year-olds and is defined as

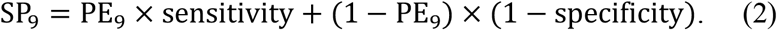

Similar to other models of CYD-TDV, our default assumption was a three-dose schedule with 100% compliance. In the event that compliance is lower, our results would be more pertinent to a scenario with a correspondingly higher coverage, as the effects of coverage and compliance are interchangeable in this way.

### Simulations of intervention impact

We performed 3,000 sets of simulations of intervention impact, with each simulation set involving one simulation with the intervention and one without. These simulation sets used the sobol function in the pomp library [32] in R [31] to evenly span a range of values of intervention coverage (10-80%), PE9 (0.1-0.9), and sensitivity (0-1) and specificity (0-1) of serological screening. Each simulation lasted for 50 years, with the intervention being introduced after the first 40 years. Every year thereafter, a proportion of nine-year-olds underwent serological screening for prior DENV exposure and were vaccinated if screening returned a positive result. Both simulations in each set were initiated with the same random number seed, which allowed us to isolate the impact of the intervention to the greatest extent possible under a stochastic, agent-based model. With each set of parameter values, we calculated the proportion of cases averted over a 10-year period as

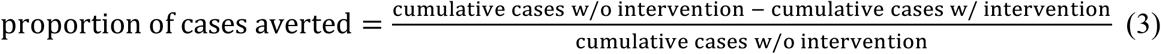

for both symptomatic and hospitalized cases. To estimate the impact of the intervention from the perspective of an individual who chose to undergo serological screening, we compared the risk of individuals from the first cohort of nine-year-olds who underwent serological screening with individuals from a comparable cohort of nine-year-olds who did not undergo serological screening. These individuals were followed for 10 years after vaccination and came from the same simulation. We calculated relative risk of symptomatic disease and hospitalization as

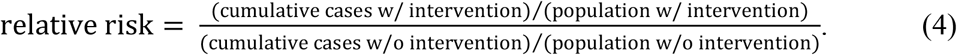

To extract average patterns from the highly stochastic outputs from 3,000 simulations of our model and to interpolate across gaps in parameter space, we summarized simulation outputs with generalized additive models, as described in Appendix S3. To assess the impact of vaccination over a longer time frame, we also evaluated effects of vaccination from both population and individual perspectives over 30 years. Results corresponding to parameter sets beyond those shown here can be explored interactively online at http://denguevaccine.crc.nd.edu.

### Identifying conditions for positive impact

Our first goal was to quantify the health impact of vaccination with CYD-TDV following serological screening under different conditions. At the population level, we made projections of the proportion of cases averted over a 10-year period, separately for symptomatic and severe cases, under a range of values of intervention coverage, PE_9_, sensitivity, and specificity. From the perspective of an individual who underwent serological screening, and vaccination in the event of a positive result, we made projections of the relative risk of experiencing a symptomatic or hospitalized case as compared to someone who forewent serological screening altogether. We examined this individual risk in aggregate and stratified by prior DENV exposure.

### Identifying conditions for cost-effectiveness

Our second goal was to understand the conditions under which vaccination with CYD-TDV following serological screening might be cost-effective. The intervention was deemed cost-effective if

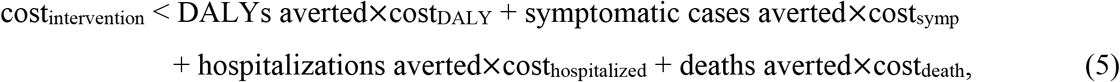

where cost_symp_ and cost_hospitalized_ reflect costs of ambulatory care and inpatient hospital care for symptomatic and hospitalized cases, respectively, and cost_death_ refers to the direct cost of death, such as burial expenses and disruption to family income. DALYs refer to disability-adjusted life years, which are years of healthy life lost to disease. We based calculations of DALYs averted on three components: symptomatic cases averted and the DALYs associated with a symptomatic case, hospitalized cases averted and the DALYs associated with a hospitalized case, and deaths averted and the average number of years of life lost for an individual in our model with a dengue-associated death. The cost of a DALY, cost_DALY_, was based on a country’s gross domestic product (GDP) per capita, in line with WHO guidance [33]. An intervention with cost_intervention_ satisfying eqn. 5 was deemed “cost-effective” when cost_DALY_ = 3 x per capita GDP and “very cost-effective” when cost_DALY_ = 1 x per capita GDP. Our assumptions about the numerical values of costs in Brazil and the Philippines are based on previous estimates used by Flasche et al. [7] and are detailed in Table 1. We applied a 3% annual discounting rate to both costs and DALYs.

**Table 1.** Assumed costs and DALYs associated with dengue illness.

| Parameter | Brazil |  | Philippines |  |
| --- | --- | --- | --- | --- |
|  | Public payer | Individual | Public Payer | Individual |
| cost<sub>symp</sub> | \$60 [36] | \$140 [5,37] | \$20 [4,38] | \$20 [4,37] |
| cost<sub>hospitalized</sub> | \$200 [36,37] | \$300 [5,37] | \$400 [5,38] | \$100 [5,37] |
| cost<sub>death</sub> | - | \$11,000 [7] | - | \$3,000 [7] |
| Per capita GDP | \$8,649.95 [39] | | \$2,951 [39] | |
| DALYs of symptomatic cases | 0.006 [40] |  |  |  |
| DALYs of hospitalized cases | 0.02 [40] |  |  |  |
| DALYs of fatal cases | 1 x years of life lost |  |  |  |

We took two approaches from the perspective of the cost of the intervention, which is defined as

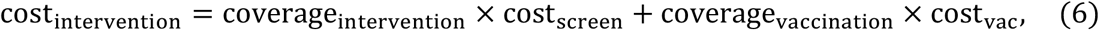

where cost_screen_ is the unit cost of serological screening and cost_vac_ is the cost of fully vaccinating a single person. Our first approach involved seeking the threshold cost of serological screening at which costs below that threshold would be cost-effective when combined with a cost_vac_ of 69 USD, which we based on pricing information from the Philippines [34] as explained in Appendix S4. Our second approach involved determining whether a fixed cost_screen_ of 10 USD (similar to a recent estimate of 9.25 USD in Vietnam [22]) would result in cost-effectiveness under three different assumptions about cost_vac_ corresponding to three, two, or one doses (69, 46, or 23 USD), assuming that any number of doses confers the same degree of protection. The possibility that fewer than three doses may confer protection against dengue has been suggested as a possibility but requires further investigation [35]. Under both approaches, we examined how cost-effectiveness varied as a function of intervention coverage, PE_9_, and the sensitivity and specificity of serological screening.

Aspects of our cost-effectiveness analysis also differed depending on the perspective of who was paying for the intervention: either a public payer (e.g., government or healthcare provider) or an individual. Health benefits in terms of cases and deaths averted differ from these population and individual perspectives, with the former being of interest to a public payer. Costs from these perspectives were differentiated in two ways. First, we monetized the direct cost of death, cost_death_, from the individual perspective as one year of productivity lost, as previously assumed by Flasche et al. [7], but we assumed no additional direct costs of fatal cases from the public payer perspective. Both perspectives considered the cost of death associated with DALYs due to premature death. Second, we assumed that ambulatory care and hospitalization costs were different for the individual and the public payer. Specific assumptions about costs from these perspectives are provided in Table 1.

## RESULTS

### Conditions for positive health impact from a population perspective

The proportion of cases averted depended on the sensitivity and specificity of serological screening in different ways for different values of PE_9_. In terms of symptomatic cases, the intervention resulted in a positive impact under nearly all combinations of parameters in all transmission settings. This was a consequence of the fact that calibration of our model to data from CYD-TDV trials resulted in estimates of the probability of symptomatic disease that decreased with each successive infection (Table 2). Thus, vaccinating more people, regardless of serostatus, resulted in more symptomatic cases averted (Fig. 1, top). In terms of hospitalizations averted, the intervention resulted in a negative impact under approximately half of the scenarios we examined. Specifically, impact was more positive in settings with higher transmission and more negative in settings with lower transmission (Fig. 1, bottom). With respect to screening properties, sensitivity was the dominant factor in high-transmission settings, and specificity was the dominant factor in low-transmission settings. For both symptomatic and hospitalized cases, relationships at lower values of PE_9_ were less smooth, due to a larger influence of stochasticity and more uncertainty in these transmission settings (Fig. S2).

**Table 2.** Parameters describing vaccine profile calibrated to CYD-TDV trial data [9]. Details of the calibration procedure are described in Appendix S2.

| Parameter | Estimate |
| --- | --- |
| Per-exposure protection from vaccination for seronegative vaccinees | 0.321 |
| Per-exposure protection from vaccination for seropositive vaccinees | 0.516 |
| Average duration of protection for seronegative vaccinees | 426 days |
| Average duration of protection for seropositive vaccinees | 258 days |
| Probability of symptoms conditional on infection (primary) | 0.405 |
| Probability of symptoms conditional on infection (secondary) | 0.339 |
| Probability of symptoms conditional on infection (post-secondary) | 0.09 |
| Probability of hospitalization conditional on symptoms (primary) | 0.074 |
| Probability of hospitalization conditional on symptoms (secondary) | 0.376 |
| Probability of hospitalization conditional on symptoms (post-secondary) | 0.101 |

**Figure 1.**
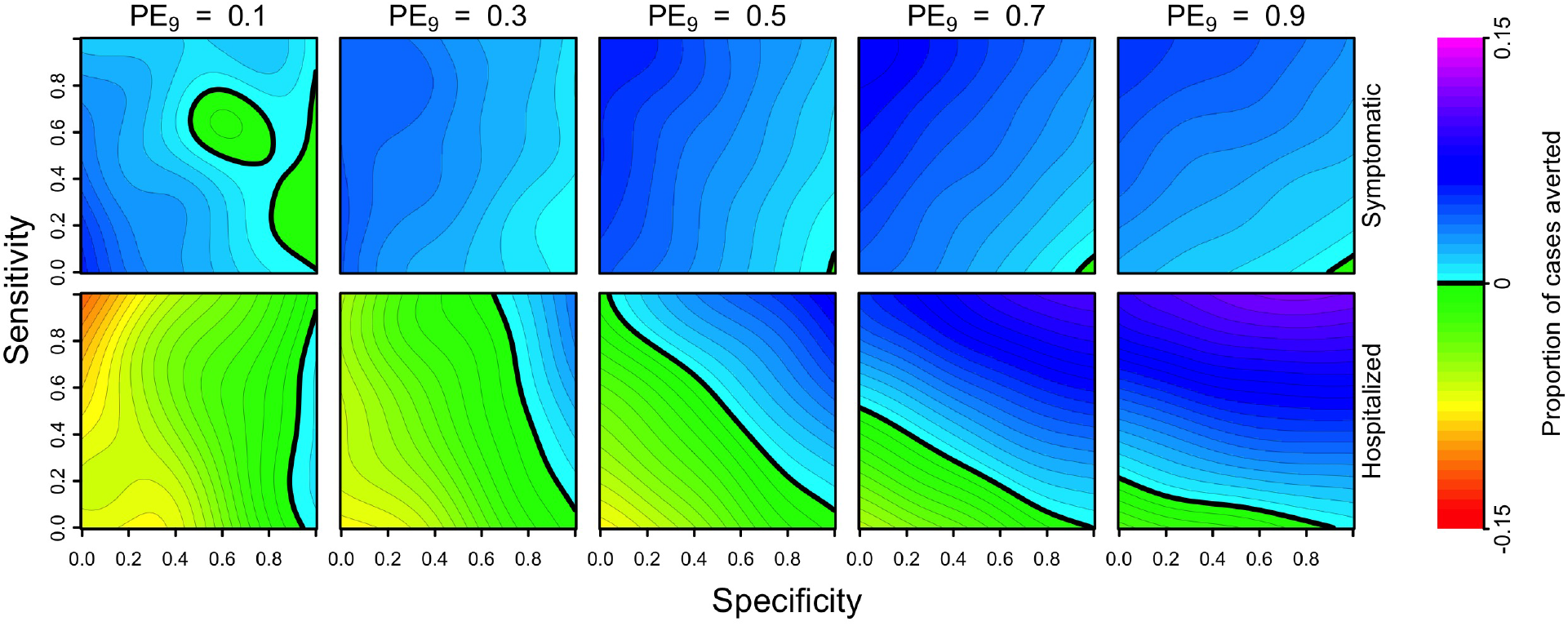
Cumulative proportion of cases averted (colors) over a 10-year period (top: symptomatic, bottom: hospitalized) as a function of the sensitivity (y-axis) and specificity (x-axis) of serological screening. Each column shows results for a given transmission setting, defined by the proportion of nine-year-olds with previous DENV exposure, PE_9_. Relationships at lower values of PE_9_ were less smooth, due to a larger influence of stochasticity and more uncertainty in these transmission settings (Fig. S2). The strategy of vaccination without screening is represented in the top-left corner of each heatmap (sensitivity = 1, specificity = 0).

The primary explanation for the positive relationship between screening sensitivity and cases averted in the highest PE_9_ setting (0.9) is that vaccination coverage depended almost exclusively on sensitivity and very little on specificity (Fig. S1). From a population perspective, achieving high coverage in a high-PE_9_ setting appeared ideal, although it also appeared that high specificity had benefits in high-transmission settings by increasing the proportion of hospitalized cases averted (11% for sensitivity = 1, specificity = 1) beyond levels achievable by high vaccination coverage alone (9% for sensitivity = 1, specificity = 0) (Fig. 1, bottom right). At low PE_9_, coverage was highest when specificity was low (Fig. S1), but that resulted in an increased number of DENV-naïve vaccinees who then went on to experience symptomatic disease and possibly hospitalization upon natural infection (Fig. 1, bottom left). Thus, public health impact was maximized at low PE_9_ when specificity was high (which minimized individual harm) and sensitivity was also high (which increased coverage among the few who should have been vaccinated).

### Conditions for positive health impact from an individual perspective

From the perspective of a nine-year-old who underwent serological screening (and, in the event of a positive result, vaccination), the relative risk of symptomatic disease was generally reduced. Given that the vaccine reduces the hazard of symptomatic disease for both seropositive and seronegative individuals, relative risk of symptomatic disease lessened as the proportion of vaccination coverage increased (Fig. 2, top). As with population-level impacts, the relative risk of hospitalization was reduced in medium- to high-transmission settings (PE_9_≥0.5) and depended on sensitivity and specificity in other settings (PE_9_<0.5) (Fig. 2, bottom).

**Figure 2.**
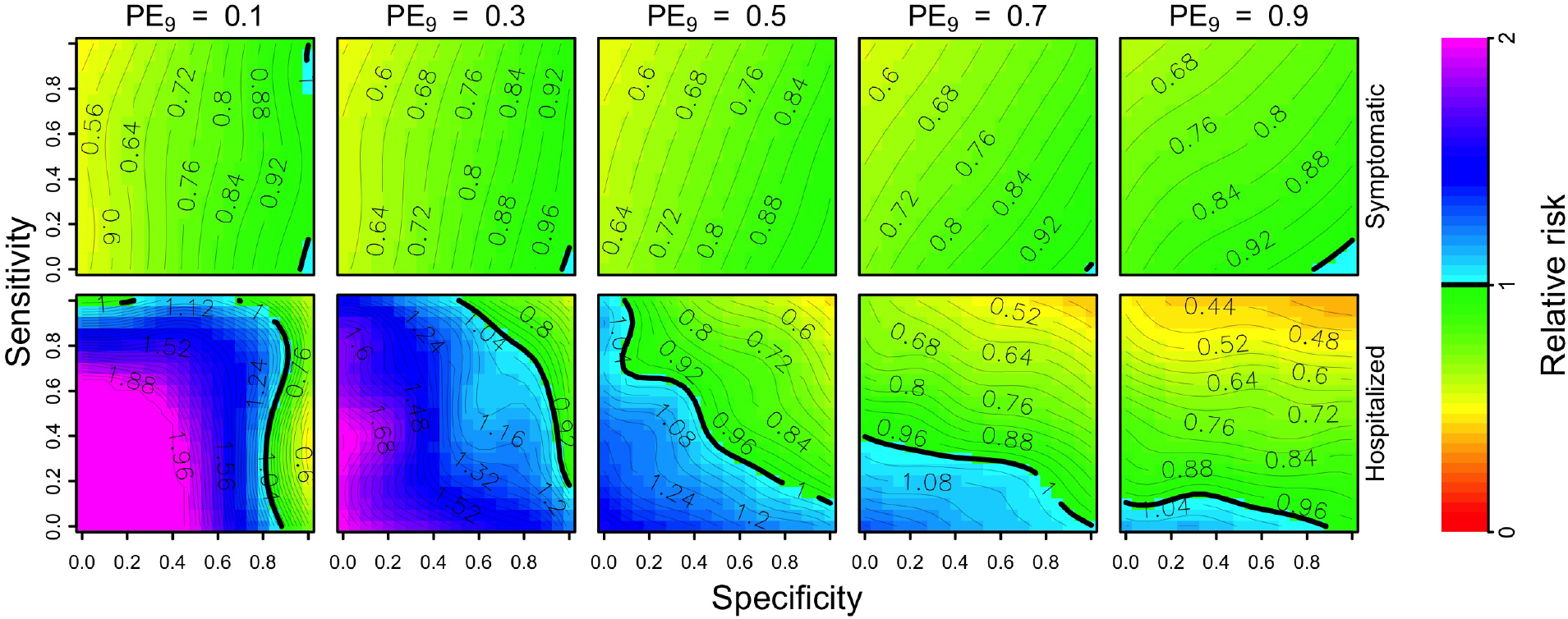
Per capita relative risk (colors) of symptomatic (top) and hospitalized (bottom) disease over a 10-year horizon in the first cohort of children who are screened (and, in the event of a positive result, vaccinated), as a function of the sensitivity (y-axis) and specificity (x-axis) of serological screening. Each column shows these results in a given transmission setting, defined by the proportion of nine-year-olds with previous DENV exposure, PE_9_. The strategy of vaccination without screening is represented in the top-left corner of each heatmap (sensitivity = 1, specificity = 0).

Under a scenario of PE_9_=0.7, relative risk of hospitalization was reduced when sensitivity was at least 0.4 or specificity was above 0.9. This reduction was mostly driven by sensitivity when specificity was below 0.8, whereas specificity modulated risk as much as sensitivity for values of specificity above 0.8. The greatest benefits occurred in high-transmission settings (PE_9_=0.9) with high sensitivity (≥0.9) and high specificity (≥0.8), in which case relative risk was as low as 0.4 (Fig. 2, bottom right). In low-transmission settings (PE_9_<0.5), relative risk of hospitalization was generally elevated, unless specificity was very high. Moreover, the reduction of risk in low-transmission settings was low, even with high specificity and sensitivity. Even though greater sensitivity reduced relative risk for an average person undergoing serological screening, from the point of view of a truly seronegative individual undergoing screening, relative risk of hospitalization was always elevated unless specificity was perfect (Fig. S3, top). In medium- to high-transmission settings (PE_9_≥0.5), relative risk was 1.1 or less for specificity values above 0.9, compared to relative risk higher than 1.3 under a scenario in which serological screening resulted in all children being vaccinated (sensitivity = 1, specificity = 0).

### Age of vaccination

Under an assumption of routine vaccination, age of vaccination modulated the population-level benefits of vaccination in terms of hospitalizations averted (Fig. S4). In higher transmission settings, vaccination at younger ages resulted in increased benefits, given that a large proportion of vaccinees had at least one infection at the time of vaccination (Fig. S4, top right). In contrast, benefits of vaccination were higher in low-transmission settings when older children were vaccinated. Vaccination in low-transmission settings appeared to have positive impacts only when routine vaccination occurred in children 15 years of age or older and specificity was high (Fig. S4, bottom left).

### Conditions for cost-effectiveness from a public payer perspective

From a public payer perspective, and assuming a cost for a full three doses of vaccine of 69 USD, our results suggest that a strategy of vaccinating seropositive nine-year-olds would be cost-effective only under limited circumstances. In simulations of medium-transmission settings (PE_9_=0.5) and with a Brazil-like scenario about costs, vaccinating seropositive nine-year-olds was cost-effective for high values of specificity (>0.8) and modest values of sensitivity (>0.3). In high-transmission settings (PE_9_ ≥ 0.7), cost-effectiveness depended on both sensitivity and specificity, with the highest thresholds for cost-effectiveness found at sensitivity and specificity above 0.8 (Fig. 3, bottom right). In a high-transmission scenario (PE=0.9), we found that the threshold cost for serological screening (i.e., the maximum cost at which the intervention could still be cost-effective) was around 45 USD. Under a Philippines-like scenario about costs, vaccinating seropositive nine-year-olds was not cost-effective under any of the scenarios that we considered (Fig. S5).

**Figure 3.**
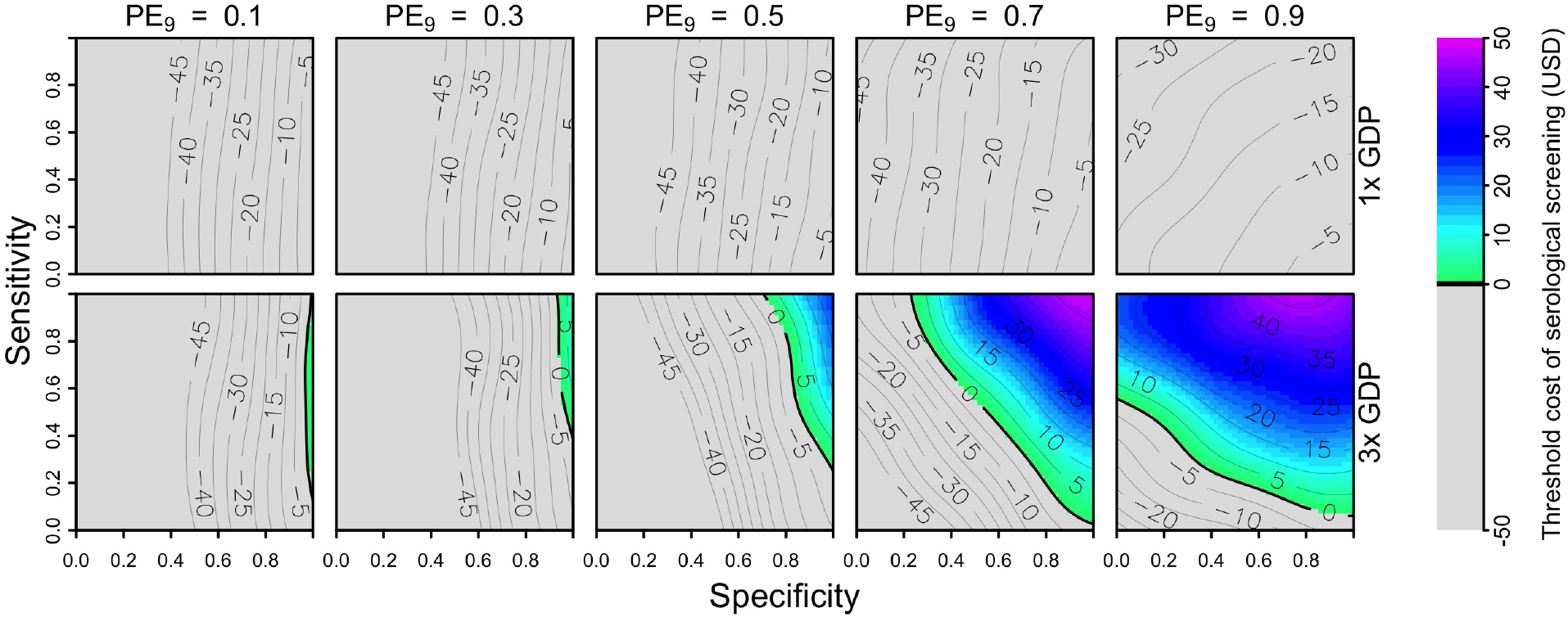
Threshold cost of serological screening from a public payer perspective, assuming a vaccination cost of 69 USD and economic assumptions from Brazil. Threshold costs are indicated by color as a function of sensitivity (y-axis), specificity (x-axis), and PE_9_ value (columns). The value of cost_DALY_ is equal to per capita GDP (8,650 USD) in the top row and three times per capita GDP in the bottom row. The strategy of vaccination without screening is represented in the top-left corner of each heatmap (sensitivity = 1, specificity = 0).

Our results showed that cost-effectiveness was possible under a somewhat broader range of parameters when we considered lower costs of the vaccine and a fixed cost of serological screening (10 USD). We found that reducing the cost of the vaccine to 46 USD (equivalent to two doses, assuming that they provide the same protection as three) had little impact on which parameter combinations (PE_9_, sensitivity, specificity) resulted in cost-effectiveness (Figs. S6 & S7). In contrast, reducing the cost of the vaccine to 23 USD (equivalent to one dose, assuming that it provides the same protection as three) resulted in cost-effectiveness in high-transmission settings (PE_9_ ≥ 0.7) under both the Brazil and Philippines scenarios about costs (Fig. 4, Fig. S8).

**Figure 4.**
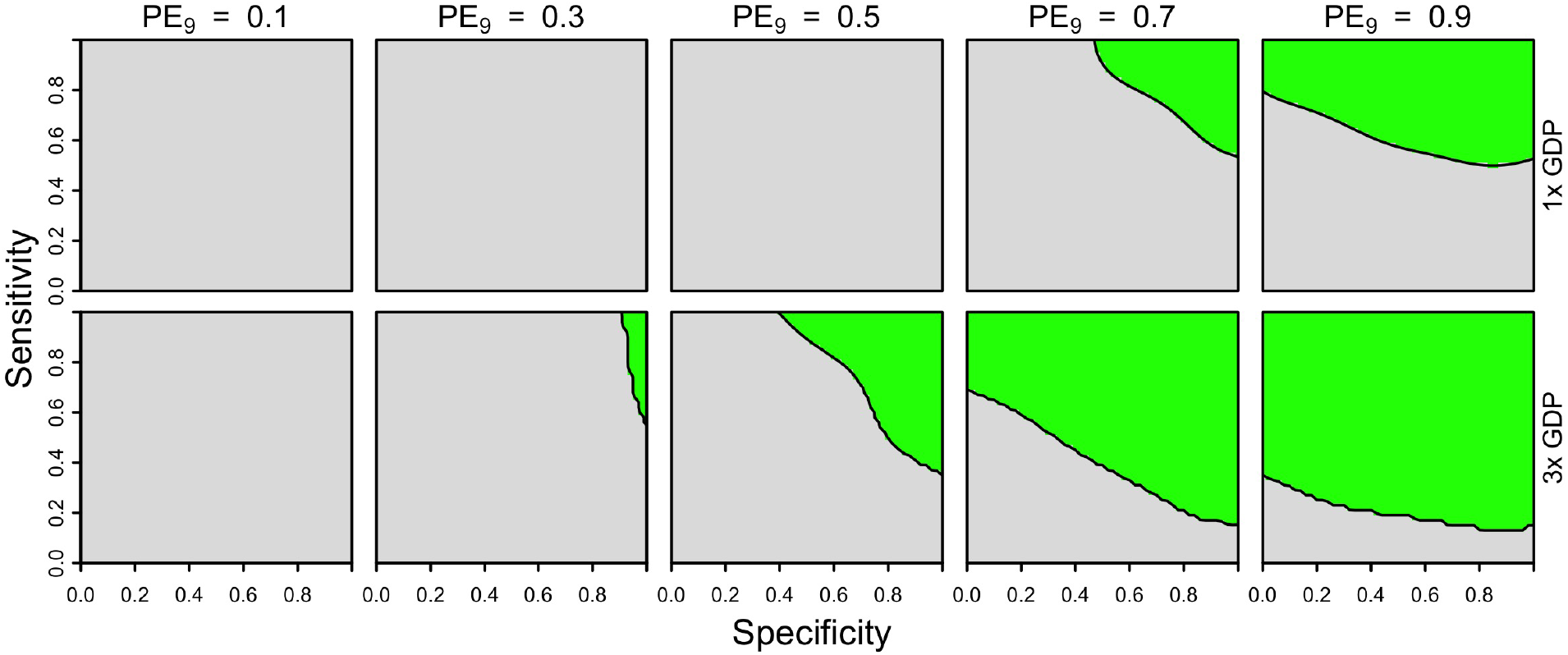
Cost-effectiveness of the intervention from a public payer perspective, assuming one dose of vaccine (23 USD) and a fixed cost of serological screening (10 USD) under Brazil-like cost assumptions. Cost-effectiveness according to eqn. 5 is shown in green as a function of sensitivity (y-axis), specificity (x-axis), and PE_9_ value (columns). The value of cost_DALY_ is equal to per capita GDP (8,650 USD) in the top row and three times per capita GDP in the bottom row. The strategy of vaccination without screening is represented in the top-left corner of each heatmap (sensitivity = 1, specificity = 0).

### Conditions for cost-effectiveness from an individual perspective

From the perspective of the parent of a nine-year-old child considering serological screening, our results suggest that the intervention would not be cost-effective in Brazil or the Philippines (Figs. 5 & S9). For both countries, low coverage (10%) had the effect of slightly increasing the threshold cost of serological screening relative to a scenario with high coverage (80%), but not enough to achieve cost-effectiveness under any parameters we considered for the Philippines (Figs. S10 & S11). This is a result of there being more to gain by an individual opting for the intervention when coverage is lower, due to lower indirect protection from others who are vaccinated. Lowering the number of doses to two (46 USD) did not improve cost-effectiveness for the Brazil-like cost scenario (Fig. S12), although lowering to one dose (23 USD) and assuming a cost of serological screening of 10 USD did (Fig. 6, bottom). Cost-effectiveness under these scenarios in moderate transmission settings (PE=0.5) depended on high sensitivity (>0.9) and moderate specificity (>0.5). In high-transmission settings (PE_9_≥0.7), cost-effectiveness was achieved for sensitivity values above 0.5 (Fig. 6, bottom). None of the scenarios that we considered were cost-effective under the Philippines-like cost scenario (Figs S13 & S14).

**Figure 5.**
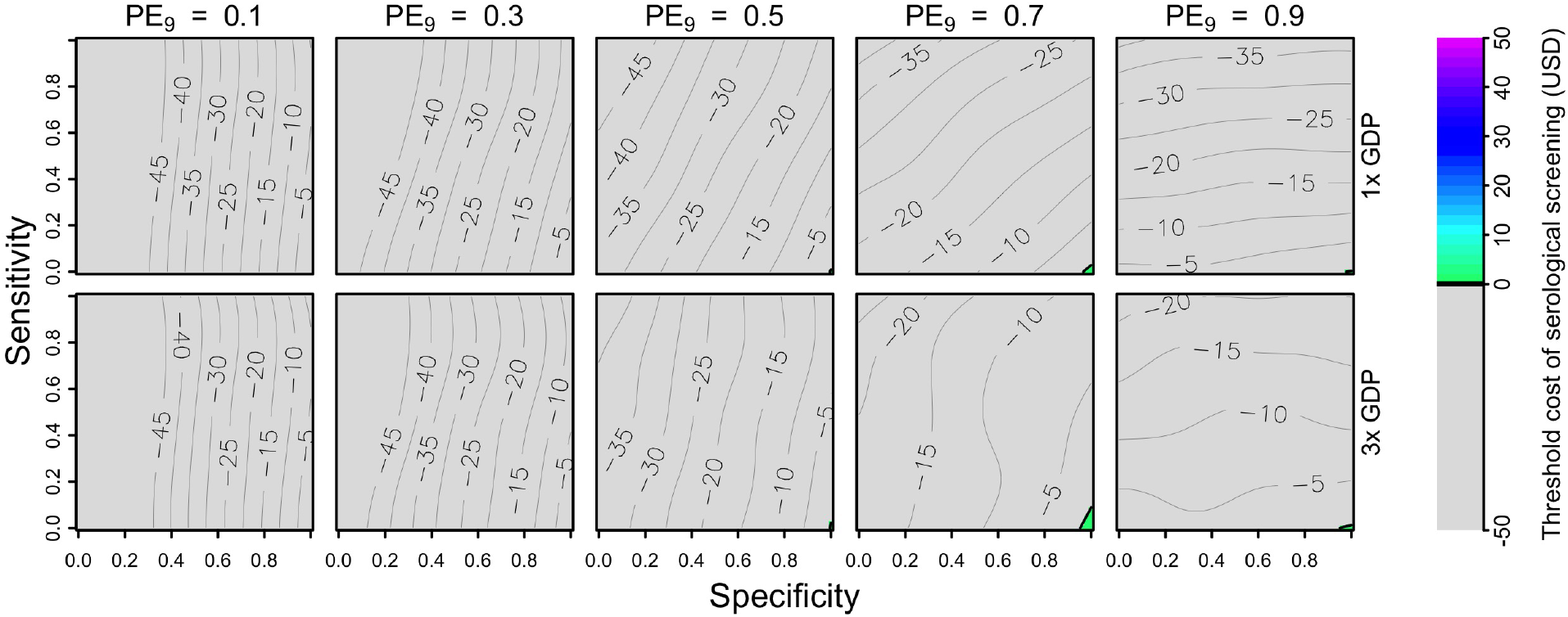
Threshold cost of serological screening from an individual perspective, assuming a vaccination cost of 69 USD and economic assumptions from Brazil. Threshold costs are indicated by color as a function of sensitivity (y-axis), specificity (x-axis), and PE_9_ value (columns). The value of cost_DALY_ is equal to per capita GDP (8,650 USD) in the top row and three times per capita GDP in the bottom row. The strategy of vaccination without screening is represented in the top-left corner of each heatmap (sensitivity = 1, specificity = 0).

**Figure 6.**
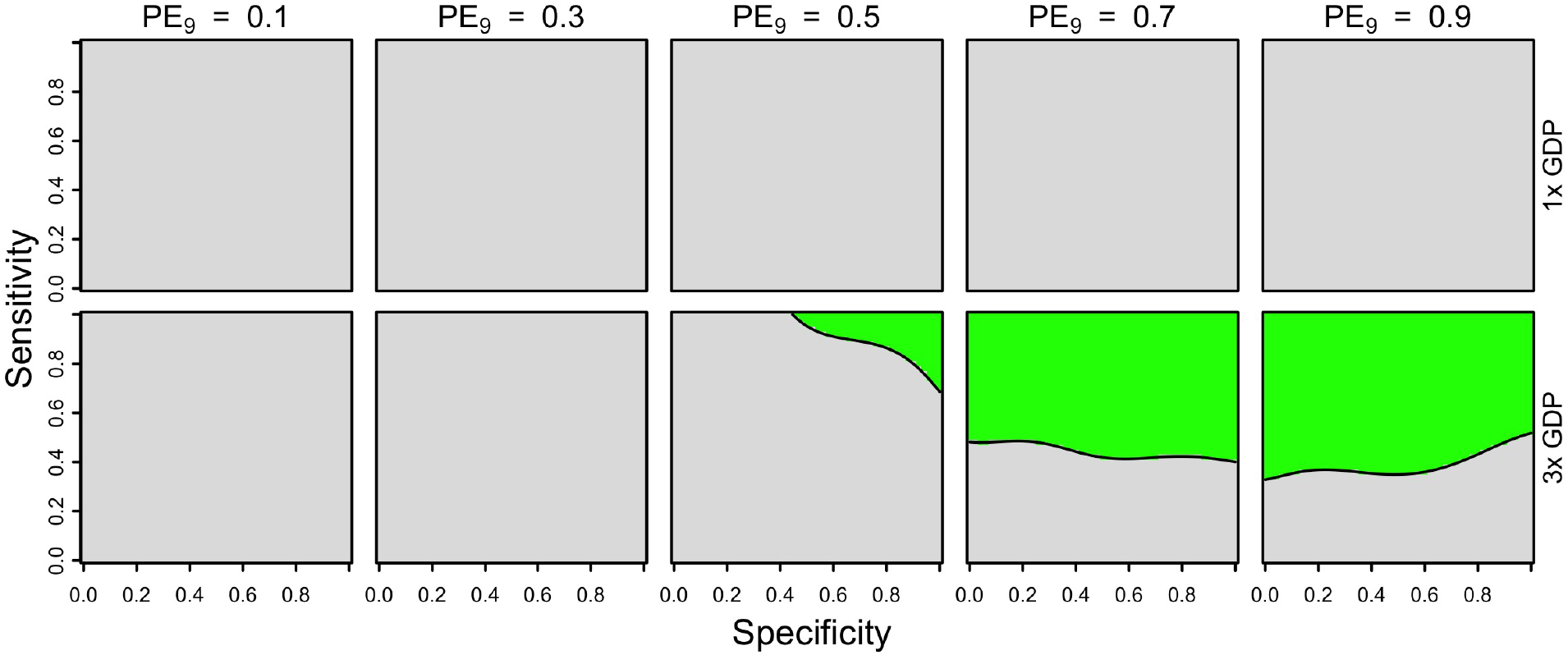
Cost-effectiveness of the intervention from an individual perspective, assuming one dose of vaccine (23 USD) and a fixed cost of serological screening (10 USD) under Brazil-like cost assumptions. Cost-effectiveness according to eqn. 5 is shown in green as a function of sensitivity (y-axis), specificity (x-axis), and PE_9_ value (columns). The value of cost_DALY_ is equal to per capita GDP (8,650 USD) in the top row and three times per capita GDP in the bottom row. The strategy of vaccination without screening is represented in the top-left corner of each heatmap (sensitivity = 1, specificity = 0).

### Health impact and cost-effectiveness over a 30-year period

Over a 30-year period, the public health impacts of the intervention were more pronounced than over a 10-year period (Figs. S15 & 1). This was true for both positive impacts in high-transmission settings and negative impacts in low-transmission settings. From an individual perspective, the magnitude of relative risk differed very little over 10-year and 30-year periods (Figs. S16 & 2). From both public health and individual perspectives, positive impacts were observed across a slightly wider range of sensitivity and specificity values (Figs. S15 & S16). Cost-effectiveness also increased from both of these perspectives, due to the fact that the cost of the intervention was the same over both time periods (Fig. S17–S24).

## DISCUSSION

Using a model consistent with seven others that informed the WHO’s initial position on CYD-TDV [7,41] but updated with the latest clinical trial data [9], we assessed the potential health impact and cost-effectiveness of the recent WHO recommendation [12] for vaccination with CYD-TDV following serological screening. In some respects, our projections were similar to previous results about vaccination without serological screening; namely, positive public health impacts in areas with high previous exposure [6,7]. In other respects, our results provide new insights on issues unique to the context of the WHO’s pre-vaccination screening recommendation.

First, our results show that high specificity is essential for reducing hospitalizations in low-transmission settings but, at the same time, leads to fewer symptomatic cases averted. The latter effect resulted from our assumption that the probability of symptomatic disease is highest in primary infections and decreases with each successive infection. Models that differ in this assumption would likely reach different conclusions about this issue. Second, our results show that sensitivity is important for achieving positive health impacts in high-transmission settings, due to the fact that higher sensitivity increases population coverage in those settings. Sensitivity appears to be less important in low-transmission settings though, from both population and individual perspectives. Third, from a public payer perspective, we conclude that cost-effectiveness is unlikely except in countries with relatively high GDP and assuming low costs of serological screening (10 USD) and vaccination (23 USD). Even then, cost-effectiveness would be limited to areas with relatively high transmission intensity and to tests with relatively high sensitivity. Fourth, conditions for cost-effectiveness from an individual perspective were more limited than from a public payer perspective. In low-transmission settings or with a low-sensitivity test in high-transmission settings, this results from the fact that the many who test negative pay to get tested but receive no health benefit as a result.

Like other modeling assessments of interventions under consideration for implementation [42–45], our study focused on offering general insights. As a consequence, we were only able to explore a relatively limited range of scenarios about vaccine roll-out. In reality, CYD-TDV could be deployed in a top-down manner by governments, purchased by individuals, or some combination thereof, given that is licensed for use in individuals ranging in age from nine to 45 years. Nevertheless, certain aspects of our analysis may offer insights about a broader range of scenarios. For example, some of our results about routine vaccination in nine-year-olds may apply under alternative scenarios if our parameter for prior exposure among nine-year-olds, PE_9_, is interpreted more broadly as prior exposure among vaccine recipients on the whole, at whatever age that might be. Such an extrapolation would appropriately mimic the level of prior exposure among vaccinees, but it may not accurately reflect transmission intensity in a population in which that level of prior exposure is achieved by a different age. At least within the 9-16 age range for routine vaccination that we considered, results from simulations involving routine vaccination in nine-year-olds appeared reasonably robust.

With respect to economic considerations, our results indicate that serological screening, and vaccination in the event of a positive result, could be cost-effective only under certain circumstances. Assuming as others have [46–48] that decisions about cost-effectiveness are made in reference to a multiplier between per capita GDP and cost_DALY_, our results predict that cost-effectiveness could be achieved only in high-transmission areas of dengue-endemic countries with a relatively high per capita GDP, such as Panamá (13,680 USD), Brazil (8,649 USD), México (8,201 USD), or Thailand (5,807 USD) [39]. In the event that CYD-TDV vaccination is recommended in a country but remains unfunded, it is likely that coverage and impact will be low, similar to varicella vaccines in Australia and Canada [49–51]. To the extent that access to CYD-TDV becomes associated with the economic means to pay for serological screening and vaccination, this could exacerbate socioeconomic disparities in dengue’s burden.

It is also important to note that our analysis of cost-effectiveness does not imply affordability. Multiple studies have shown that interventions that have been deemed “very cost-effective” have nonetheless not been implemented in low- and middle-income countries due to a variety of factors, such as implications for spending on competing public health priorities [52–54]. Another approach to estimating cost_DALY_ is to refer to incremental cost-effectiveness ratios (ICERs) from other interventions that could be displaced by CYD-TDV, such as vaccines against rotavirus and human papillomavirus. These interventions have been shown to be very cost-effective in settings comparable to Brazil and the Philippines, with ICERs below 2,000 [55,56]. Based on our results, none of the scenarios that we considered would result in cost-effectiveness of CYD-TDV comparable to these interventions, given that that would have required cost-effectiveness to be achieved with cost_DALY_ < 2,000 USD.

Although our analysis provides an indication of desirable characteristics of assays for serological screening, there is not yet an assay available that is simultaneously rapid, point-of-care, low-cost, and both highly sensitive and specific [57]. Neutralization assays, for example, are reasonably accurate but expensive and time-consuming, whereas assays such as IgG ELISAs are faster and relatively inexpensive, but often far less accurate [20]. Given the tradeoffs between the sensitivity and specificity of any assay, our results suggest that priority should be placed on maximizing specificity. Doing so would minimize the potential risks associated with vaccination of DENV-naïve individuals misclassified by an imperfectly-specific assay as having been previously exposed. Achieving high specificity in determining DENV serostatus is complicated by numerous sources of cross-reactivity, including prior exposure to or vaccination against Japanese encephalitis, West Nile, yellow fever, or Zika viruses [16]. Because these factors affecting cross-reactivity are population-specific, any assay used to inform vaccination with CYD-TDV should be calibrated to results from a highly specific assay (e.g., plaque-reduction neutralization tests) in a given population to maximize specificity [58]. Inevitably though, maximizing specificity will come at the cost of decreased sensitivity [21] and, as we have shown, reduced population-level benefits. By considering the full range of possible sensitivities and specificities, our results offer a quantitative basis for assessing the potential impact and cost-effectiveness of any existing or future assay.

In theory, a highly effective, tetravalent dengue vaccine could have a substantial impact on reducing dengue’s considerable burden, but that goal remains elusive for numerous reasons [59]. In the absence of a single intervention that is highly effective across a wide range of contexts, interest continues to grow in determining how to best combine multiple interventions in ways that are appropriate for a given local context [60]. Making that determination has become increasingly challenging due to nuanced, yet highly consequential, issues associated with use of CYD-TDV. Mathematical modeling analyses offer important capabilities for addressing this challenge due to their ability to weigh complex tradeoffs among intervention properties, as demonstrated here with respect to the sensitivity and specificity of serological screening, prior DENV exposure among vaccinees, and intervention coverage and cost. In addition, by considering both individual and population perspectives, our analysis provides information that could be informative for discourse about difficult ethical considerations surrounding the use of CYD-TDV [61].

## SUPPORTING INFORMATION

### Appendix S1. Process for achieving a desired value of PE_9_ in model simulations

To afford the model the flexibility to achieve a range of transmission intensities, as defined by PE_9_, we developed a statistical emulator of PE_9_ to interpolate simulation results across gaps in parameter space of three unknown model parameters (rate at which DENV is seeded into the population, mosquito infectiousness, adult female mosquito emergence rate). To do so, we generated 10^3^ combinations of these three parameters using the sobol function in the pomp library [32] in R [31]. This function generates points that maximize distance between them within a prescribed range of values for each parameter (DENV seeding rate: 8×10^−6^−2×10^−4^; mosquito infectiousness: 0-1; adult female mosquito emergence rate: 0-3). After simulating 40 years of transmission with a given set of those three parameters, we retrieved PE_9_ from all such simulations and fitted a generalized additive model of PE_9_ with independent smooth terms for each of the three parameters (R^2^ = 0.98). In subsequent simulations focused on vaccination impact, we obtained a set of the three unknown model parameters consistent with a target value of PE_9_ by repeatedly drawing sets of the three parameters until we obtained one that was associated with a value of PE_9_ within one percent of the target PE_9_ value.

### Appendix S2. Model calibration of vaccine profile to clinical trial results

We estimated parameters related to vaccine profile based on the latest estimates of vaccine efficacy and hazard ratios from a case-cohort study including the CYD15, CYD15, CYD23, and CYD57 studies [9]. We compared our model’s outputs under a given vaccine profile to outputs reported by Sridhar et al. following imputation of missing baseline serostatus using the super learner for targeted minimum loss-based estimation (TMLE) approach. We set up a virtual trial in our model to resemble the combined sample size of the trials. Our virtual trial represents logistics and procedures from field trials, as described elsewhere [28]. To match the initial conditions of the CYD-TDV trials, we adjusted the age-specific proportion of negative serostatus at baseline (24% overall) of the trial participants in the model to match estimates based on the TMLE approach. Assuming a constant force of infection, *λ*, before the trial, the proportion of seronegatives in each age group from age *i* to age *j* was calculated as

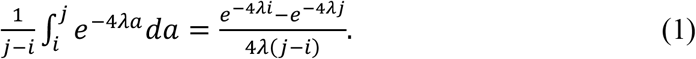

We accounted for the sensitivity and specificity of the TMLE when compared to the NS-1 titers at month 13 by re-adjusting our estimates of the proportion of seronegatives in such a way that the tested proportion of seronegatives matched the TMLE results. After this re-adjustment, we tested each individual enrolled in our virtual trial with a sensitivity of 94% and specificity of 79.1% to obtain the estimated proportion of seronegatives.

Four components of the trial were compared to the model outputs: incidence of virologically-confirmed dengue infection with symptomatic disease within the first 25 months, incidence of virologically-confirmed dengue infection with hospitalization after 60 months, vaccine efficacy estimates in the first 25 months, and the hazard ratio of hospitalization after 60 months. We calculated vaccine efficacy and hazard ratios using a Cox-regression model based on time-to-event data. The basis of our calibration method was to maximize the goodness of fit of the simulated incidence, vaccine efficacy, and hazard ratios as compared to the trial data. We measured the goodness of fit with the likelihood of the model parameters given the trial data. For the incidence of symptomatic disease and hospitalization, we assumed a binomial distribution among those eligible for each of those outcomes (DENV-infected and DENV-infected with symptomatic disease, respectively). For estimates of vaccine efficacy and hazard ratios, we assumed that the likelihood of our model’s parameters followed a normal distribution with mean and standard deviation following estimates from Sridhar et al. [9]. For each combination of parameters, the goodness of fit was estimated as the sum of the log-likelihoods.

To select the parameter values with the highest likelihood, we used a particle filter that resembles a sequential importance resampling algorithm [29]. We varied a total of 10 parameters as defined in Table 2 in the main text. A set of 2,000 combinations of parameters was proposed in the initial step of the particle filter. We used the gam function from the ‘mgcv’ [30] library in R [31] to predict the likelihood of each of the 2,000 particles and found the top 10% particles with the highest likelihood estimates. To proceed from one round of the particle filter to the next, we used a multivariate normal distribution to sample 2,000 new sets of parameters from the top 10% selected with the gam function. We performed a total of five iterations of this algorithm and selected the parameters with the highest likelihood for subsequent simulations. The calibrated model resembles the vaccine efficacy for symptomatic dengue disease (25M) and the hazard ratios of hospitalization by age-group and baseline serostatus reasonably well (Fig. S2.1).

**Fig. S2.1.**
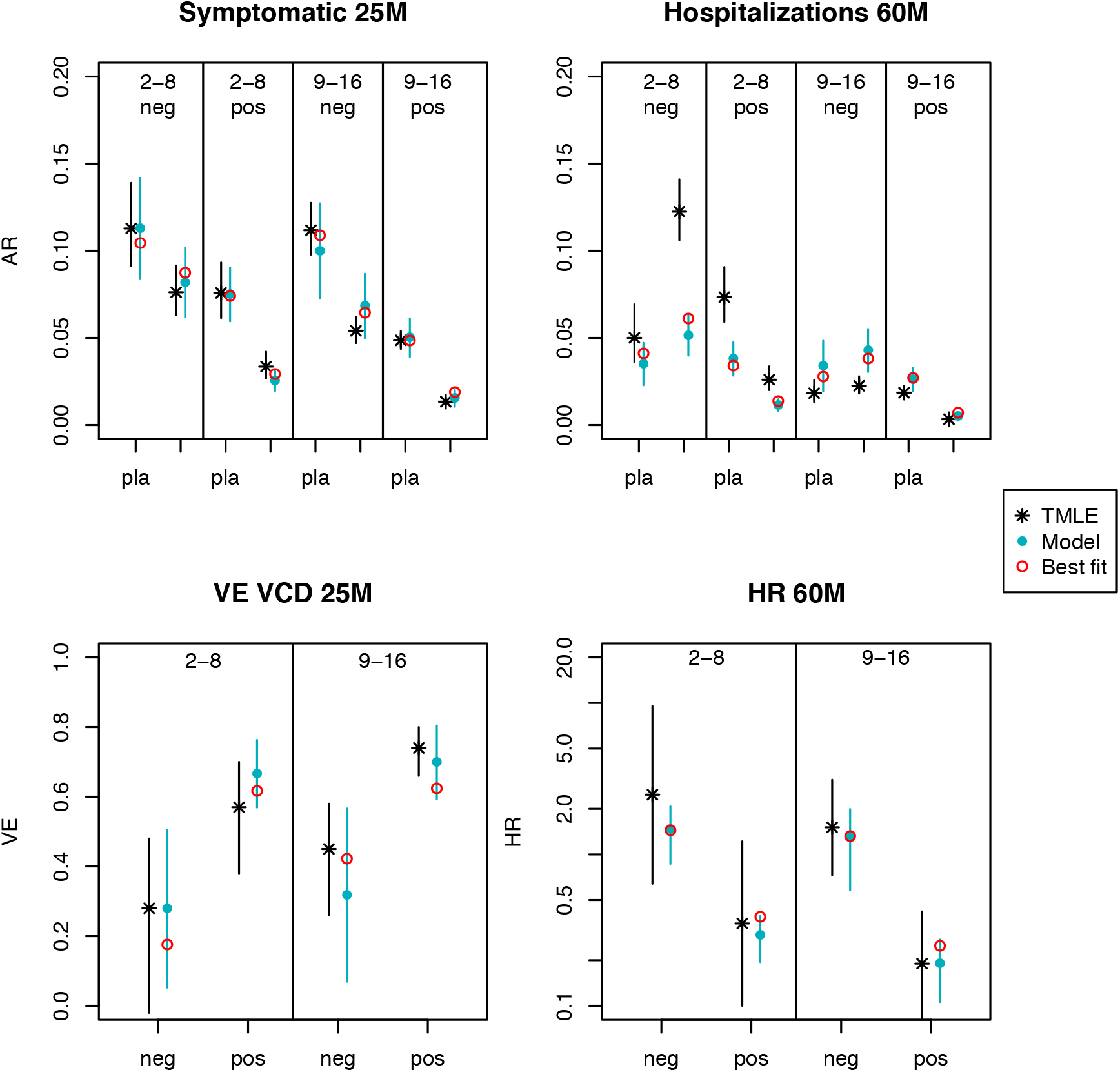
Fit of the calibrated model of vaccine profile to recent estimates of vaccine protection as described by Sridhar et al. [9]. Black dots show estimates from Sridhar et al., with black lines representing their reported 95% confidence intervals. Blue dots show the mean calibrated model response for the best 10% of particles from the last calibration step and the 95% confidence interval is indicated by the blue lines. Red dots represent the best model fit. The top left panel shows the attack rates for virologically-confirmed symptomatic dengue disease within the first 25 months of the trial. The top right panel shows the attack rates for virologically-confirmed hospitalizations due to dengue disease. The bottom left panel shows vaccine efficacy estimates for symptomatic dengue disease. The bottom right panel shows the hazard ratios following 60 months of data collection.

### Appendix S3. Generalized additive model of vaccination impact projections

Despite efforts to minimize stochastic differences between paired simulations under a given parameter set, the proportions of cases averted resulting from our simulations were relatively noisy. This is due to the dynamics of the model, which are characterized by large epidemics in some years separated by low levels of incidence during inter-epidemic periods [23]. Thus, even small differences in the sequence of random number draws (due to differences in infection outcomes associated with protective effects of vaccination) can lead otherwise similar pairs of simulations to diverge in their behavior over time. Even so, there were clear patterns in the central tendency of the proportion of cases averted as a function of the parameters varied across the 3,000 parameter sets that we examined. To extract pattern from noise, we developed a statistical emulator of the proportion of cases averted as a function of four parameters described in the previous paragraph using the gam function from the ‘mgcv’ [31] library in R [30], which fits a generalized additive model to the data. Values of the proportion of cases averted from this emulator were likewise used in calculations of cost-effectiveness.

### Appendix S4. Estimate of the price of Dengvaxia^®^ in the Philippines

In 2016, the Philippines government paid a total of P3.5 billion to vaccinate a total of 1,077,623 9-year-old public-school students [34]. We assumed that this cost allowed for three doses of vaccine plus the cost of administrating it. Hence, the unit price of a fully vaccinated person was around P3,247. This corresponded to 69.3 USD in 2016, which we rounded to 69 USD. This cost can be recalculated and updated in our analyses using the web application available online.

**Figure S1.**
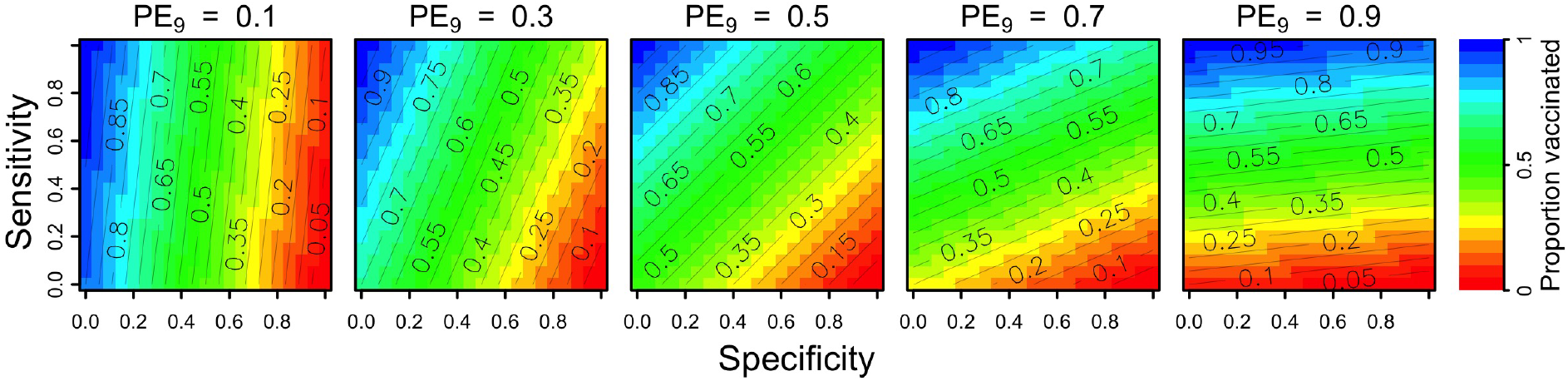
Relationship between the proportion of nine-year-olds with previous DENV exposure (columns) and the proportion who screen positive and receive vaccination (colors). This relationship depends on the sensitivity (y-axis) and specificity (x-axis) of serological screening.

**Figure S2.**
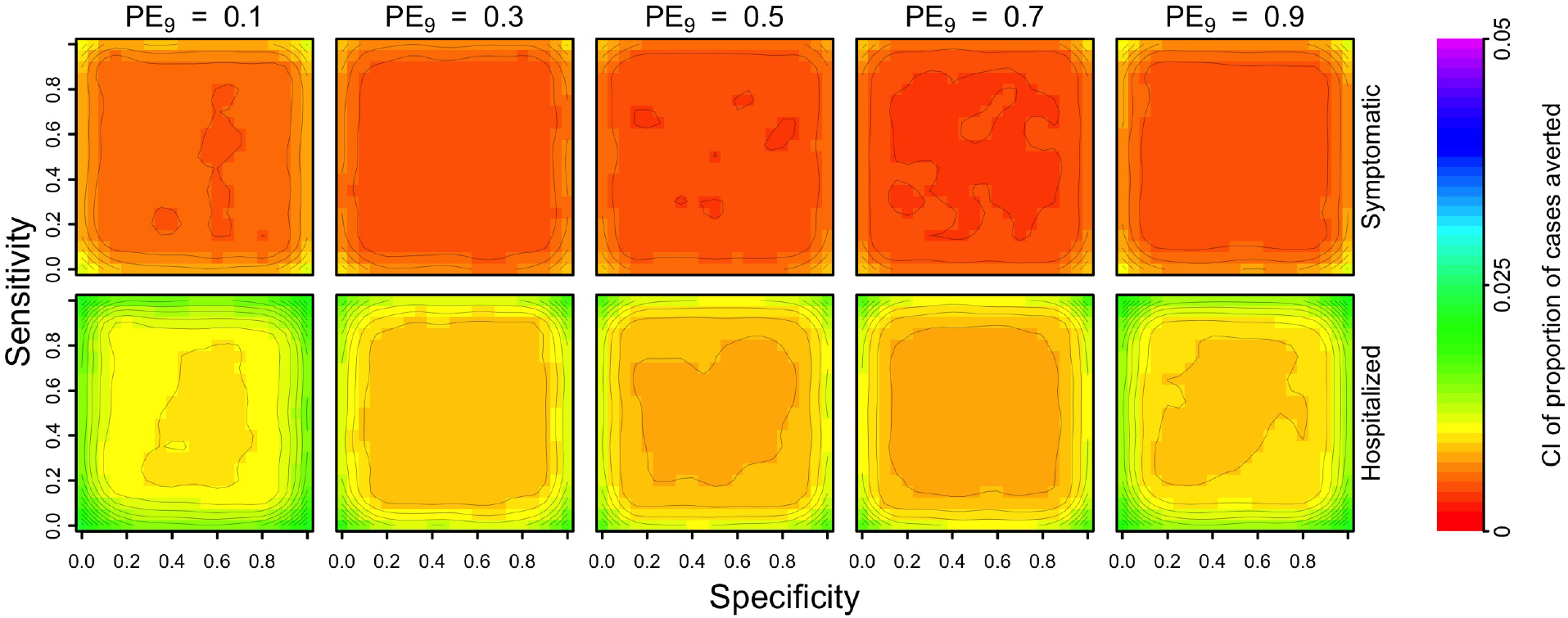
Width of the confidence interval of the cumulative proportion of cases averted over a 30-year period (top row: symptomatic, bottom row: hospitalized) as a function of the sensitivity (y-axis) and specificity (x-axis) of serological screening. Each column shows these results in a given transmission setting, defined by PE_9_.

**Figure S3.**
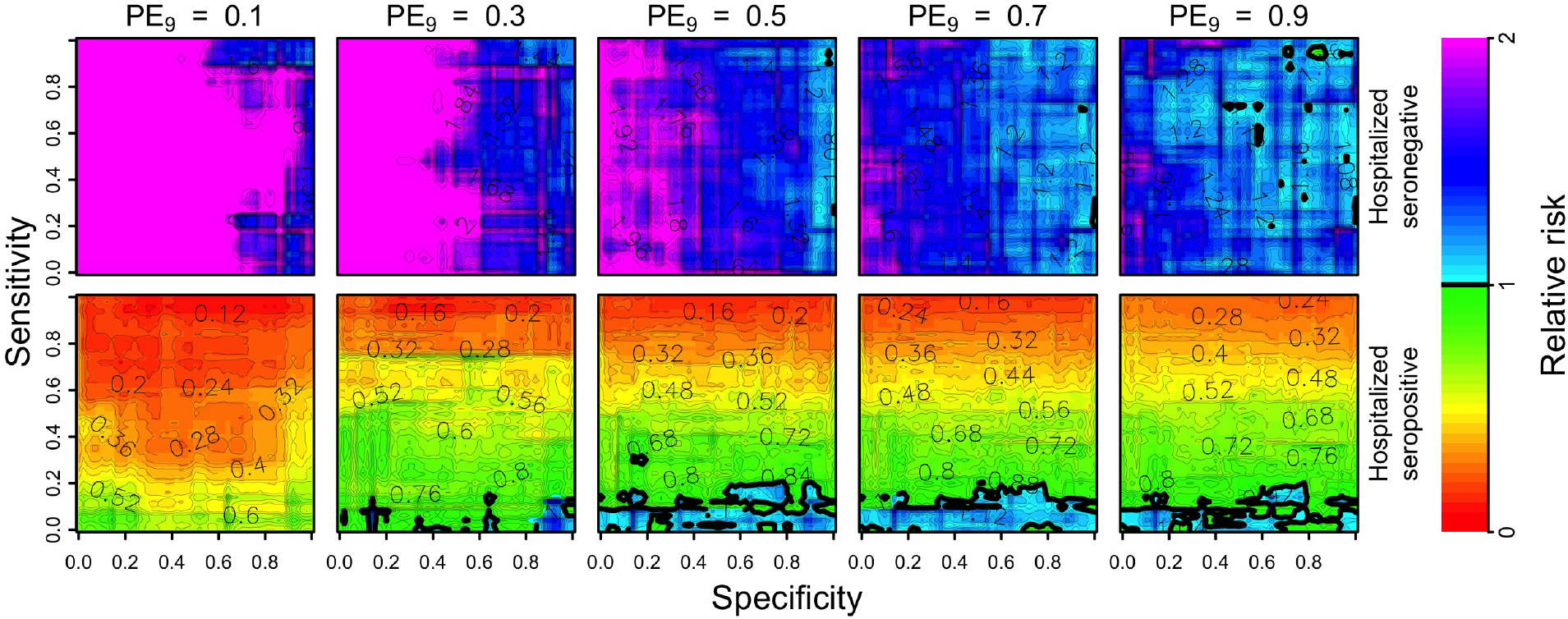
Per capita relative risk (colors) of hospitalization for individuals seronegative (top) and seropositive (bottom) over a 10-year horizon in the first cohort of individuals eligible for vaccination after a positive result from serological screening, as a function of the sensitivity (y-axis) and specificity (x-axis) of serological screening. Each column shows these results in a given transmission setting, defined by the proportion of nine-year-olds with previous DENV exposure, PE_9_.

**Figure S4.**
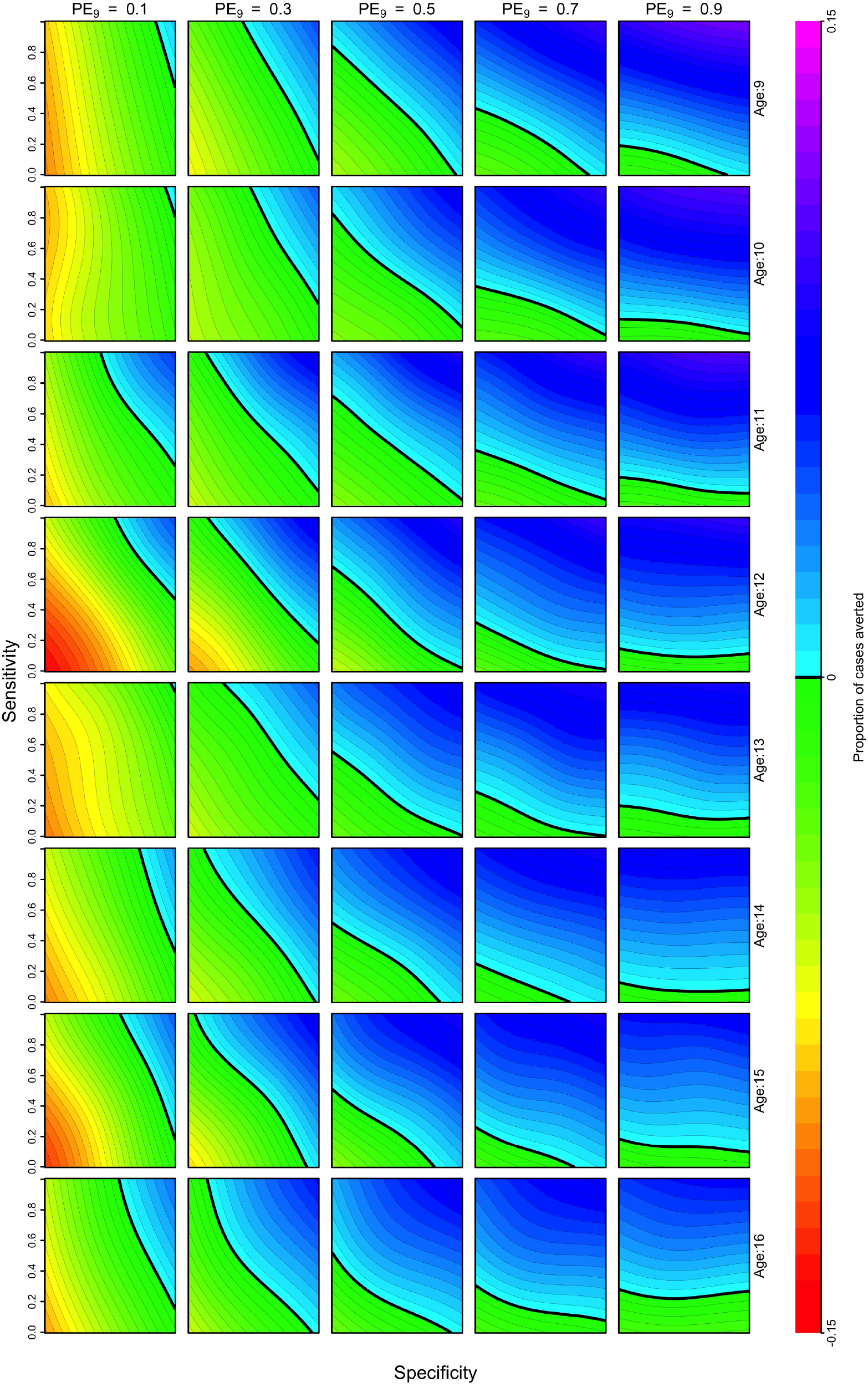
Cumulative proportion of cases averted over a 10-year period for different ages of routine vaccination.

**Figure S5.**
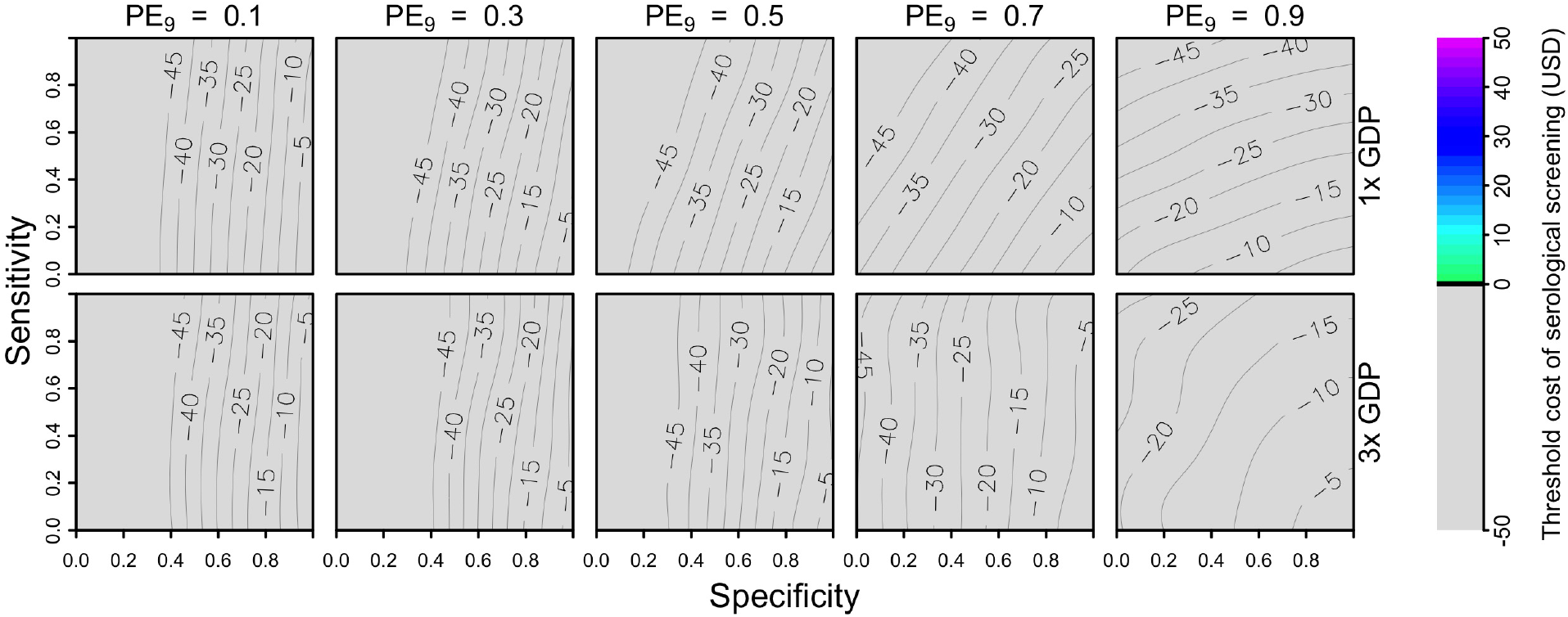
Threshold cost of serological screening from a public payer perspective over a 10-year period, assuming a vaccination cost of 69 USD and economic assumptions from the Philippines. Threshold costs are indicated by color as a function of sensitivity (y-axis), specificity (x-axis), and PE_9_ value (columns). The value of cost_DALY_ is equal to per capita GDP (2,951 USD) in the top row and three times per capita GDP in the bottom row.

**Figure S6.**
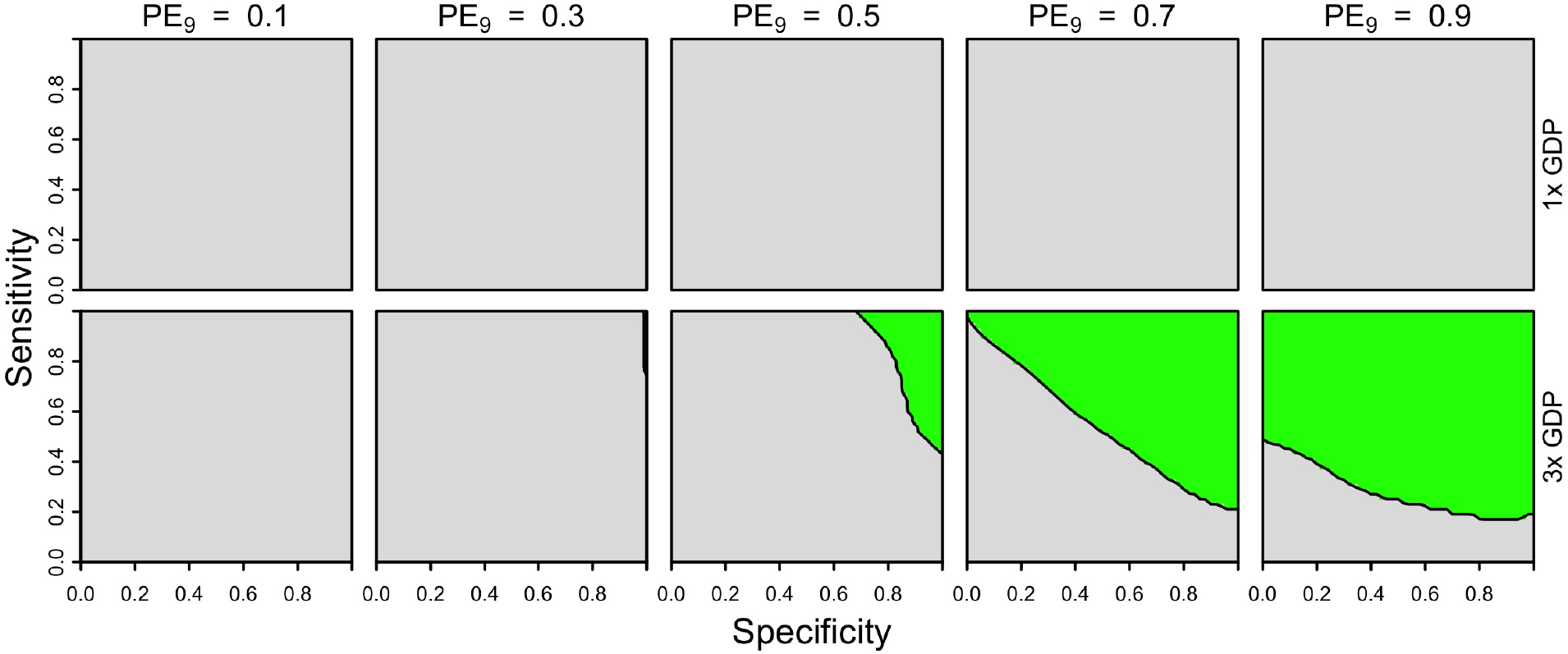
Cost-effectiveness of the intervention over a 10-year period from a public payer perspective, assuming two doses of vaccine (46 USD) and a fixed cost of serological screening (10 USD) under Brazil-like cost assumptions. Cost-effectiveness according to eqn. 5 is shown in green as a function of sensitivity (y-axis), specificity (x-axis), and PE_9_ value (columns). The value of cost_DALY_ is equal to per capita GDP (8,650 USD) in the top row and three times per capita GDP in the bottom row.

**Figure S7.**
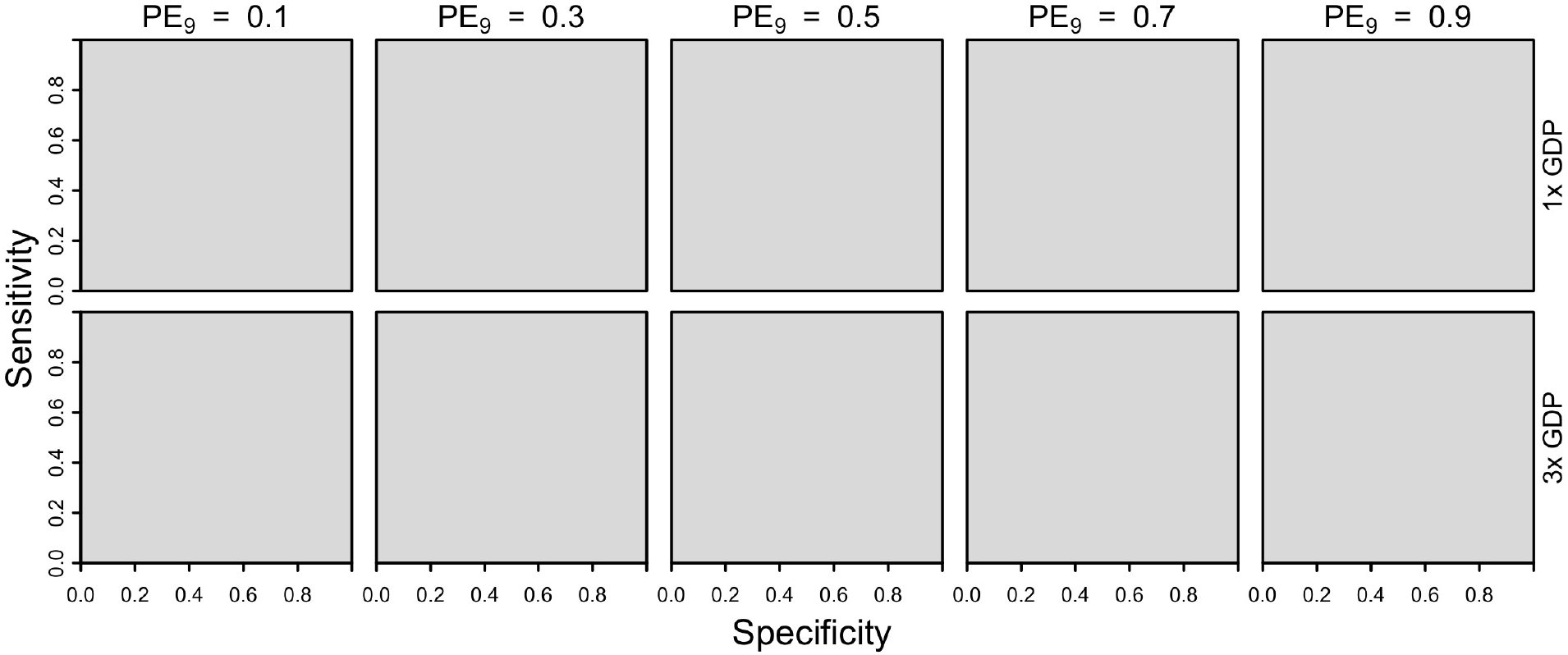
Cost-effectiveness of the intervention over a 10-year period from a public payer perspective, assuming two doses of vaccine (46 USD) and a fixed cost of serological screening (10 USD) under Philippines-like cost assumptions. Cost-effectiveness according to eqn. 5 is shown in green as a function of sensitivity (y-axis), specificity (x-axis), and PE_9_ value (columns). The value of cost_DALY_ is equal to per capita GDP (2,951 USD) in the top row and three times per capita GDP in the bottom row.

**Figure S8.**
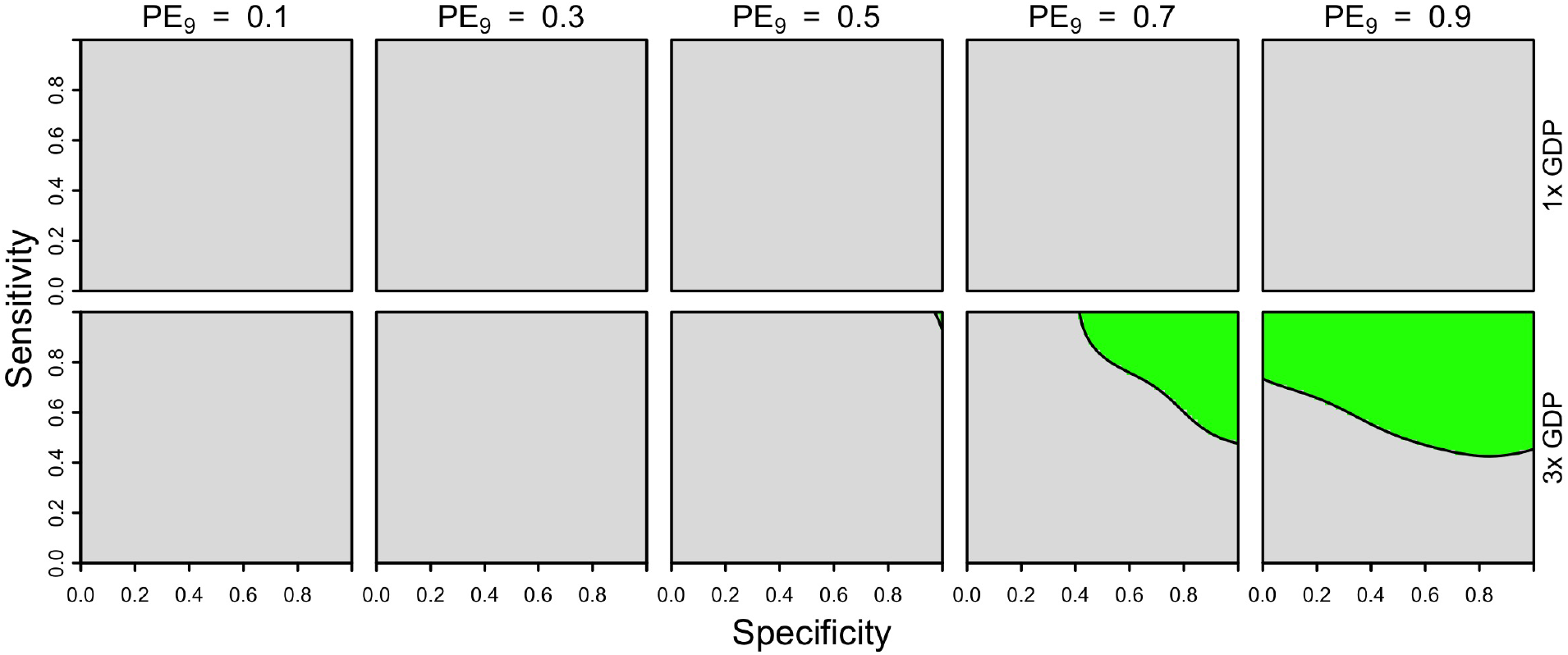
Cost-effectiveness of the intervention over a 10-year period from a public payer perspective, assuming one dose of vaccine (23 USD) and a fixed cost of serological screening (10 USD) under Philippines-like cost assumptions. Cost-effectiveness according to eqn. 5 is shown in green as a function of sensitivity (y-axis), specificity (x-axis), and PE_9_ value (columns). The value of cost_DALY_ is equal to per capita GDP (2,951 USD) in the top row and three times per capita GDP in the bottom row.

**Figure S9.**
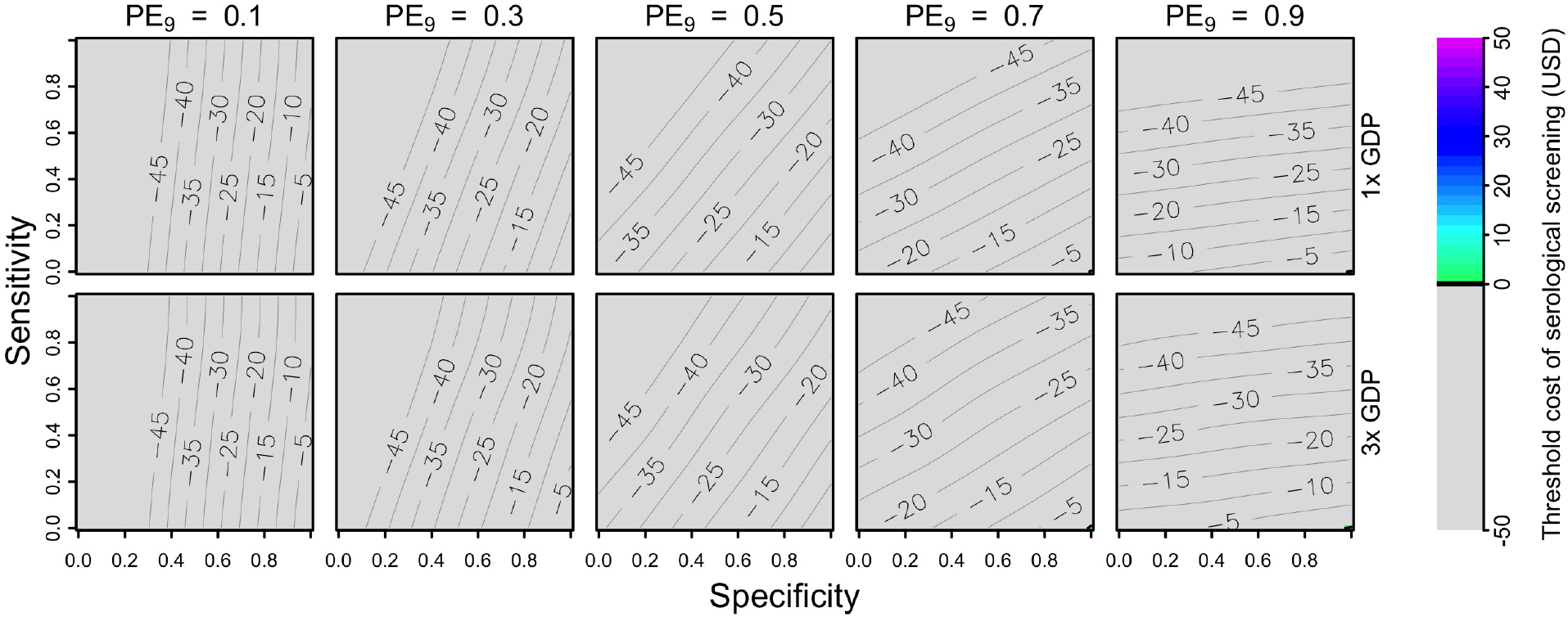
Threshold cost of serological screening from an individual perspective over a 10-year period, assuming a vaccination cost of 69 USD and economic assumptions from the Philippines. Threshold costs are indicated by color as a function of sensitivity (y-axis), specificity (x-axis), and PE_9_ value (columns). The value of cost_DALY_ is equal to per capita GDP (2,951 USD) in the top row and three times per capita GDP in the bottom row.

**Figure S10.**
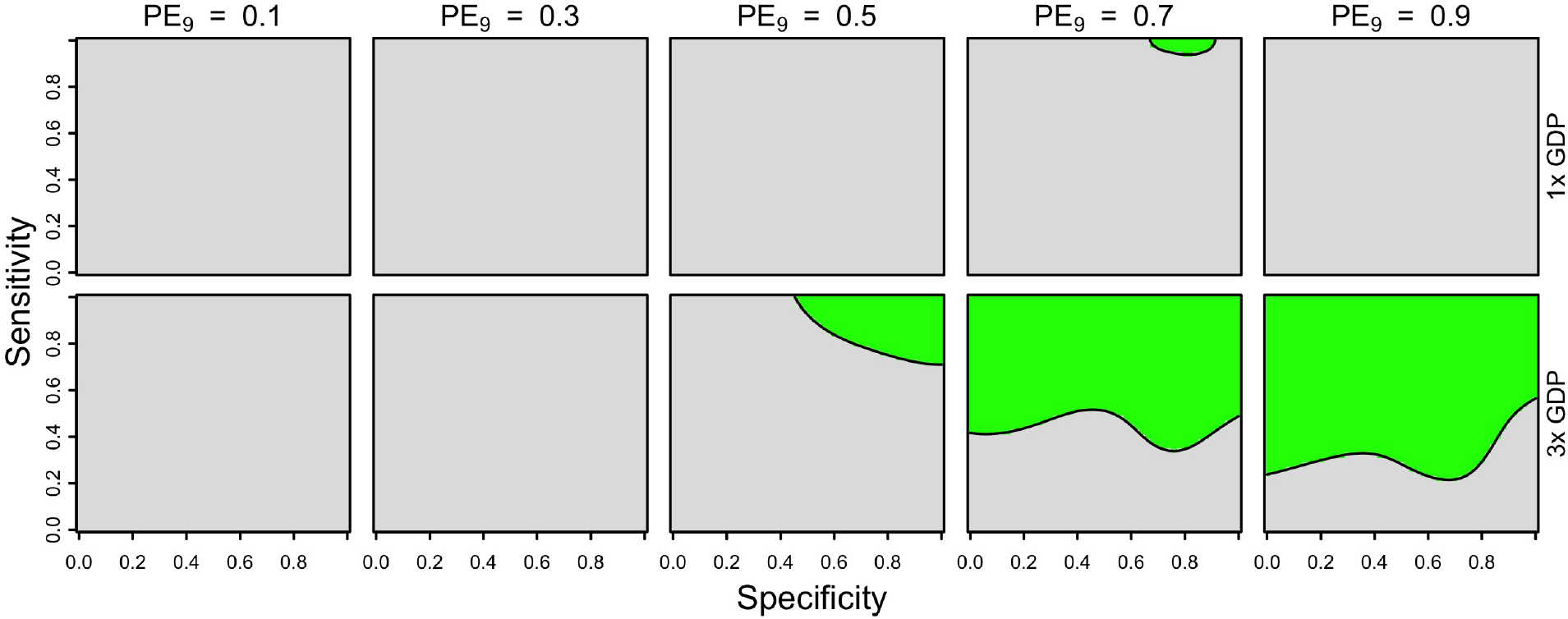
Cost-effectiveness of the intervention from an individual perspective at 10% coverage, assuming one dose of vaccine (23 USD) and a fixed cost of serological screening (10 USD) under Brazil-like cost assumptions. Cost-effectiveness according to eqn. 5 is shown in green as a function of sensitivity (y-axis), specificity (x-axis), and PE_9_ value (columns). The value of cost_DALY_ is equal to per capita GDP (8,650 USD) in the top row and three times per capita GDP in the bottom row.

**Figure S11.**
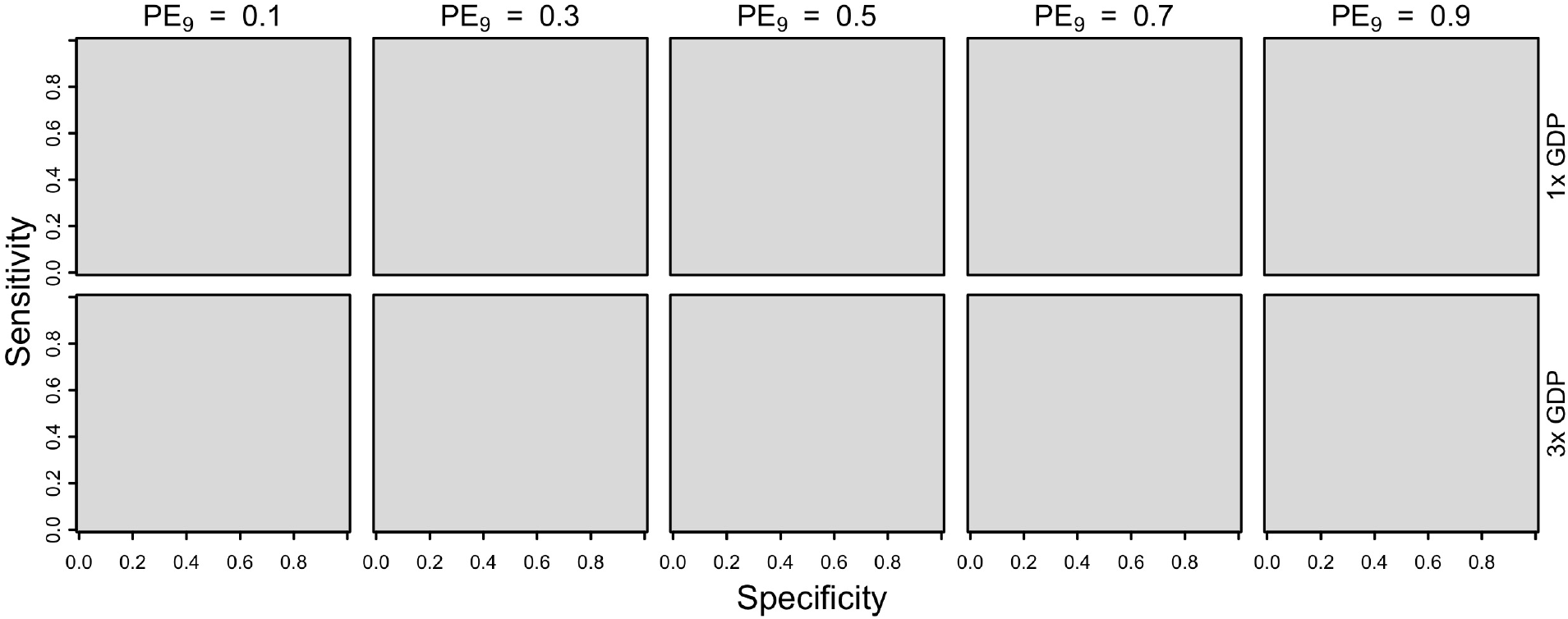
Cost-effectiveness of the intervention from an individual perspective at 10% coverage, assuming one dose of vaccine (23 USD) and a fixed cost of serological screening (10 USD) under Philippines-like cost assumptions. Cost-effectiveness according to eqn. 5 is shown in green as a function of sensitivity (y-axis), specificity (x-axis), and PE_9_ value (columns). The value of cost_DALY_ is equal to per capita GDP (2,951 USD) in the top row and three times per capita GDP in the bottom row.

**Figure S12.**
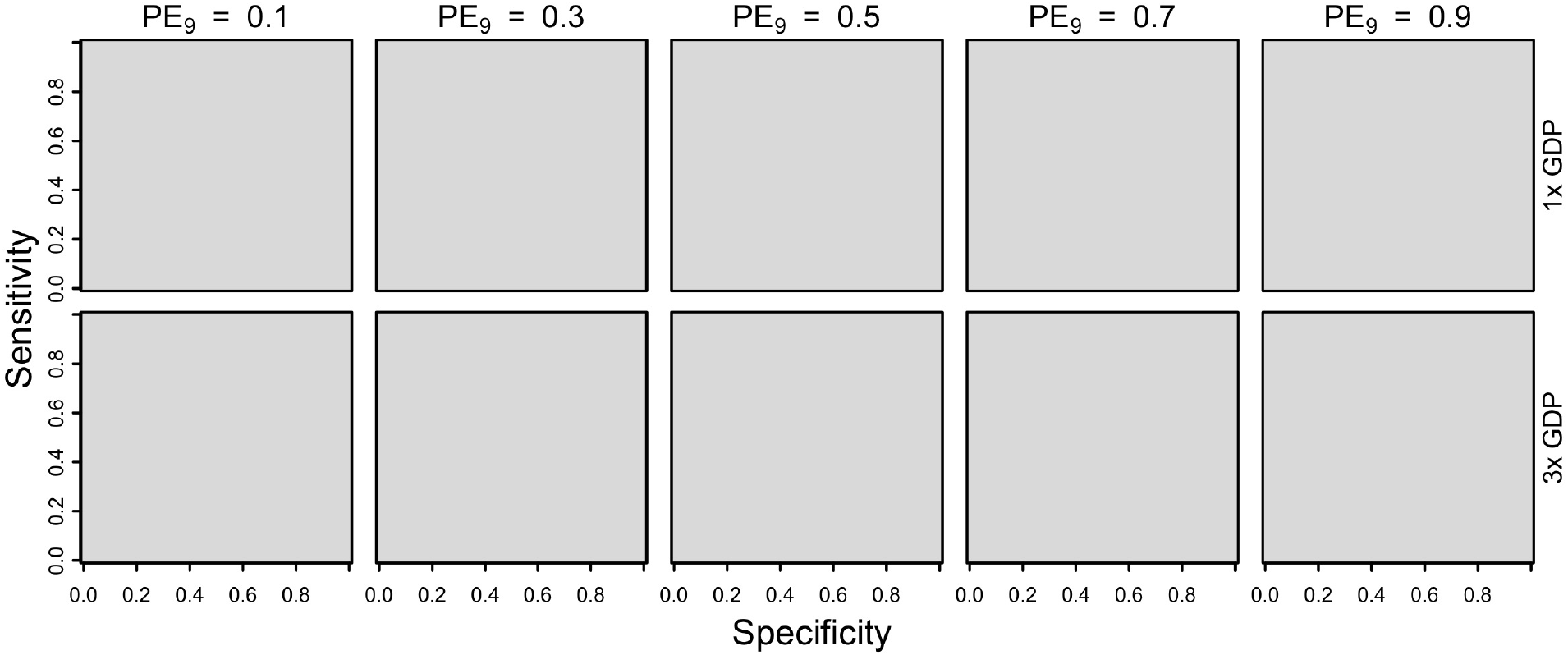
Cost-effectiveness of the intervention over a 10-year period from an individual perspective, assuming two doses of vaccine (46 USD) and a fixed cost of serological screening (10 USD) under Brazil-like cost assumptions. Cost-effectiveness according to eqn. 5 is shown in green as a function of sensitivity (y-axis), specificity (x-axis), and PE_9_ value (columns). The value of cost_DALY_ is equal to per capita GDP (8,650 USD) in the top row and three times per capita GDP in the bottom row.

**Figure 13.**
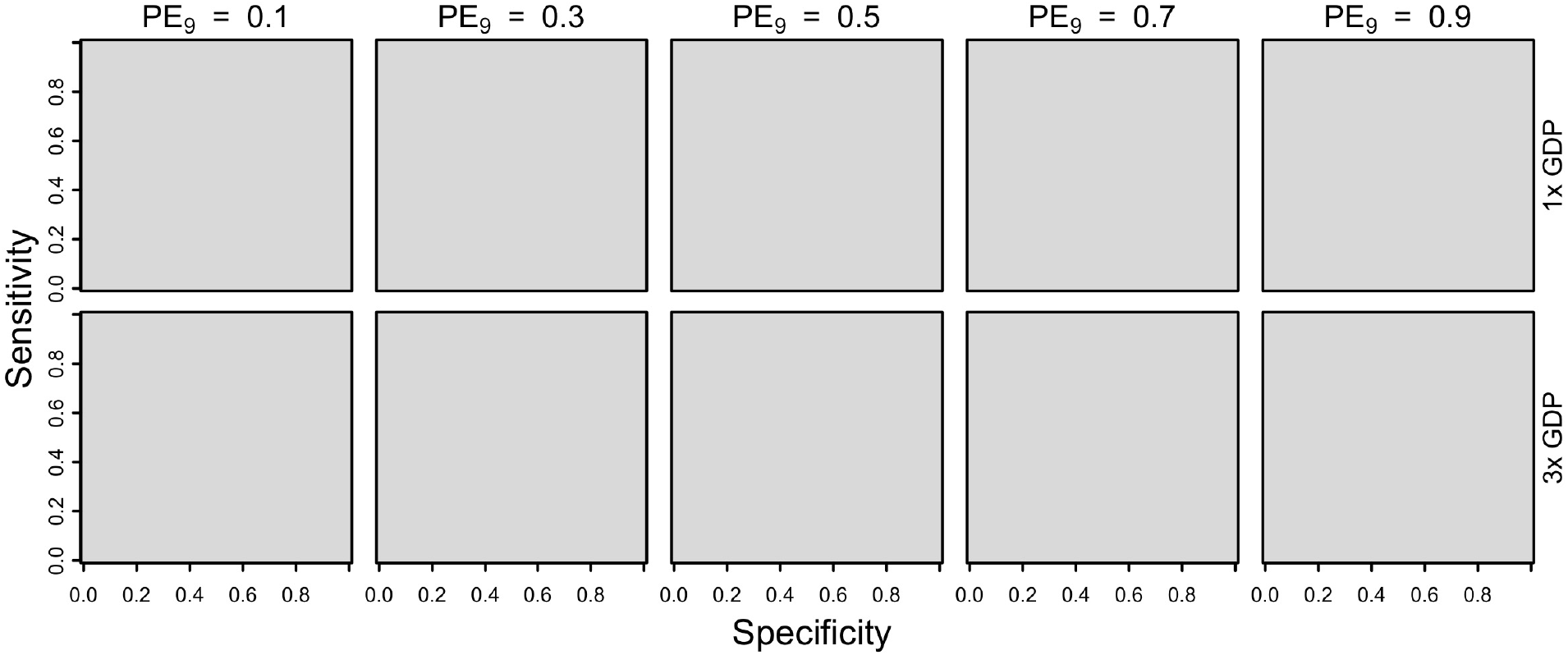
Cost-effectiveness of the intervention over a 10-year period from an individual perspective, assuming two doses of vaccine (46 USD) and a fixed cost of serological screening (10 USD) under Philippines-like cost assumptions. Cost-effectiveness according to eqn. 5 is shown in green as a function of sensitivity (y-axis), specificity (x-axis), and PE_9_ value (columns). The value of cost_DALY_ is equal to per capita GDP (2,951 USD) in the top row and three times per capita GDP in the bottom row.

**Figure S14.**
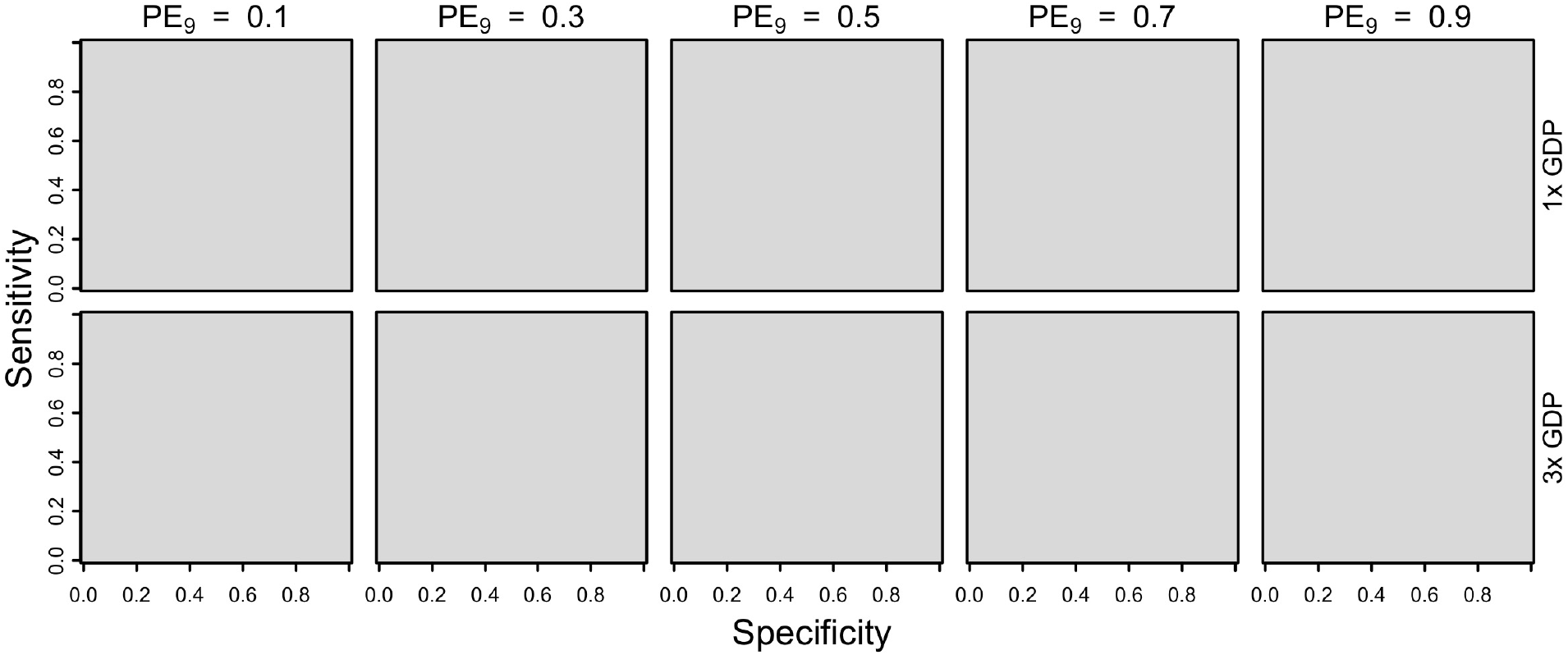
Cost-effectiveness of the intervention over a 10-year period from an individual perspective, assuming one dose of vaccine (23 USD) and a fixed cost of serological screening (10 USD) under Philippines-like cost assumptions. Cost-effectiveness according to eqn. 5 is shown in green as a function of sensitivity (y-axis), specificity (x-axis), and PE_9_ value (columns). The value of cost_DALY_ is equal to per capita GDP (2,951 USD) in the top row and three times per capita GDP in the bottom row.

**Figure S15.**
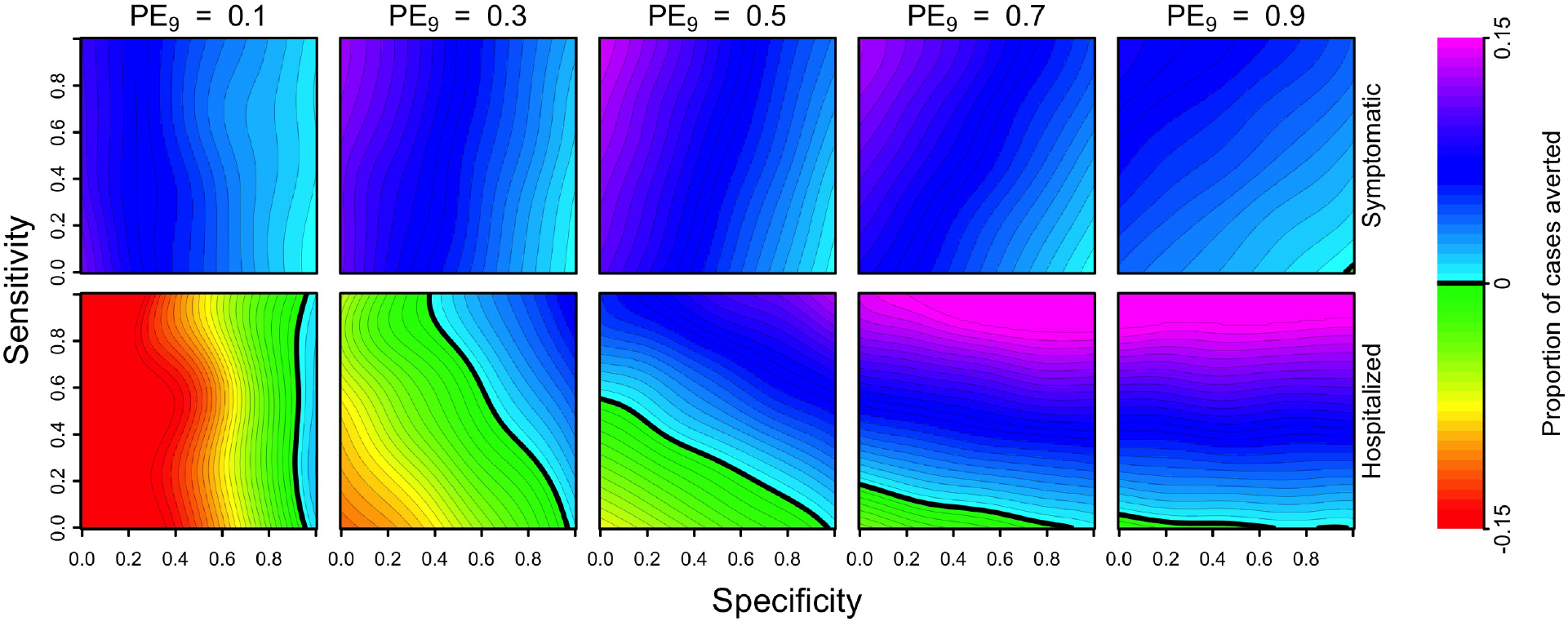
Cumulative proportion of cases averted (colors) over a 30-year period (top: symptomatic, bottom: hospitalized) as a function of the sensitivity (y-axis) and specificity (x-axis) of serological screening. Each column shows results for a given transmission setting, defined by the proportion of nine-year-olds with previous DENV exposure, PE_9_.

**Figure S16.**
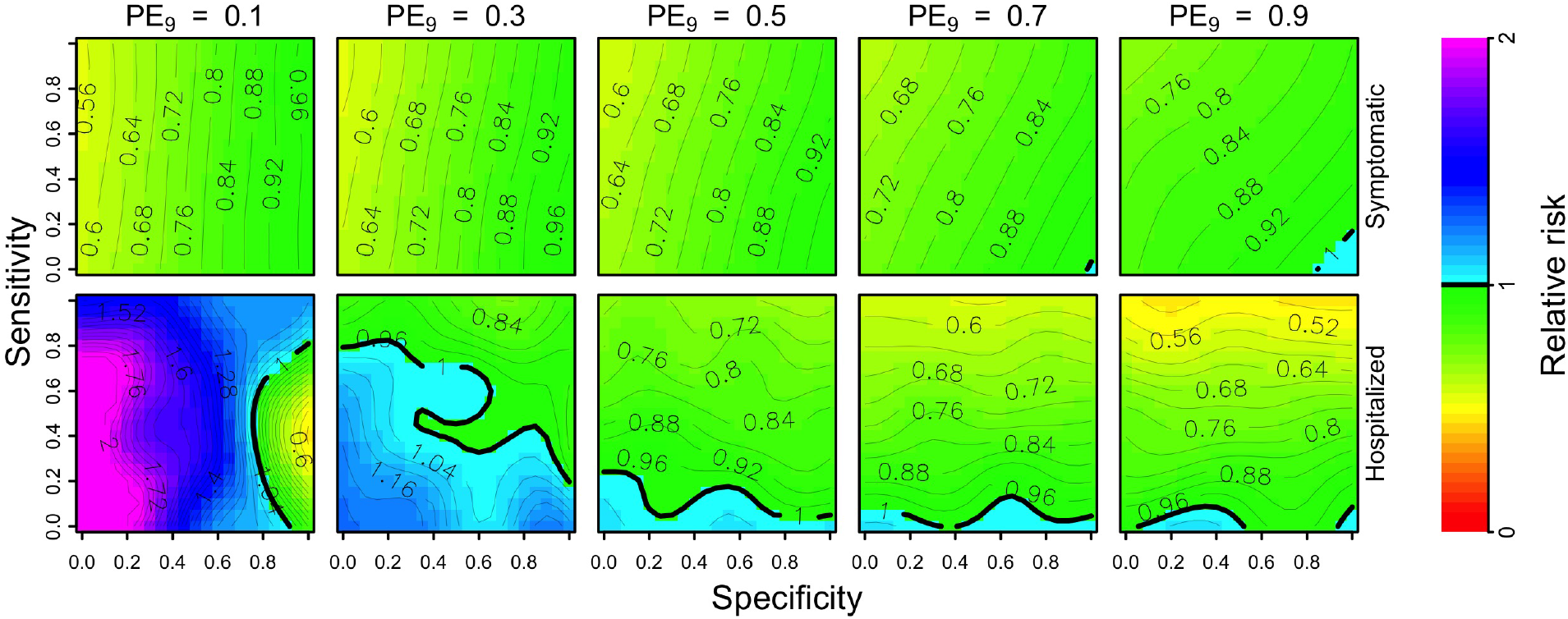
Per capita relative risk (colors) of symptomatic (top) and hospitalized (bottom) disease over a 30-year horizon in the first cohort eligible for vaccination after serological screening with a positive result, as a function of the sensitivity (y-axis) and specificity (x-axis) of serological screening. Each column shows these results in a given transmission setting, defined by the proportion of nine-year-olds with previous DENV exposure, PE_9_.

**Figure S17.**
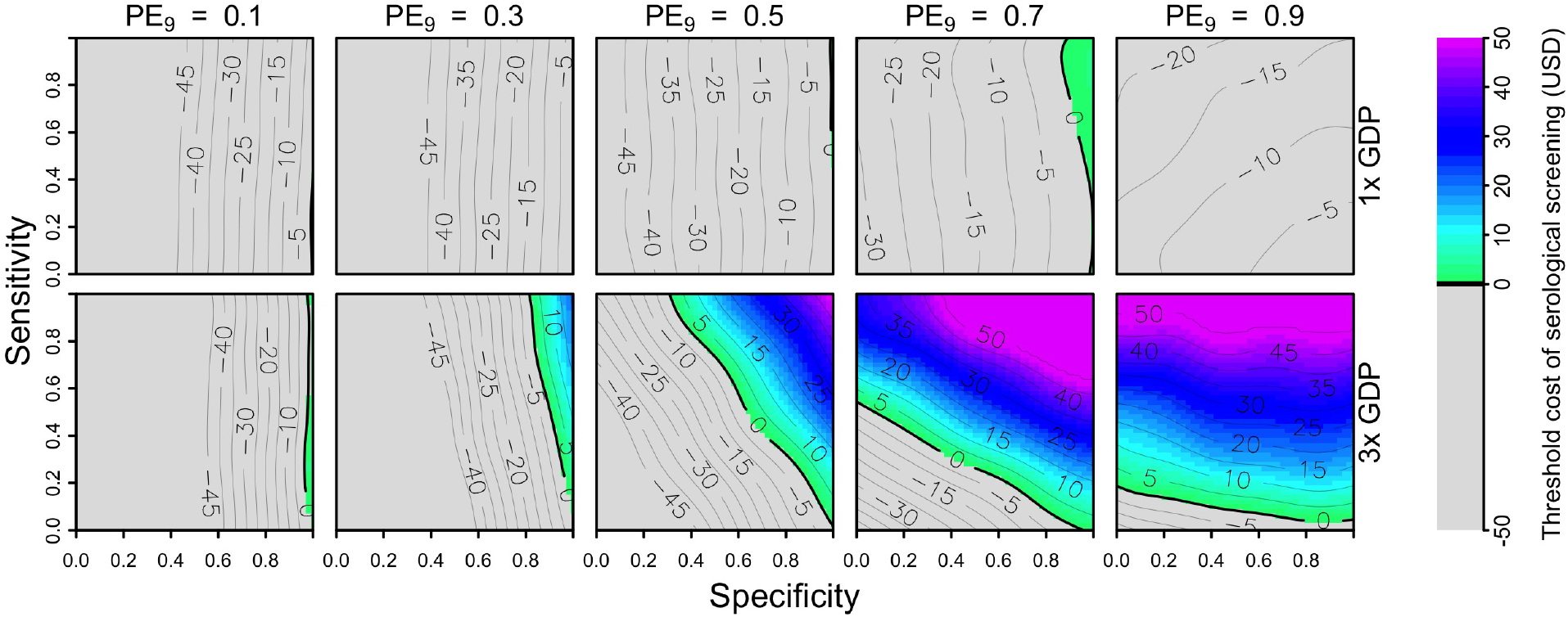
Threshold cost of serological screening from a public payer perspective over a 30-year period, assuming a vaccination cost of 69 USD and economic assumptions from Brazil. Threshold costs are indicated by color as a function of sensitivity (y-axis), specificity (x-axis), and PE_9_ value (columns). The value of cost_DALY_ is equal to per capita GDP (8,650 USD) in the top row and three times per capita GDP in the bottom row.

**Figure S18.**
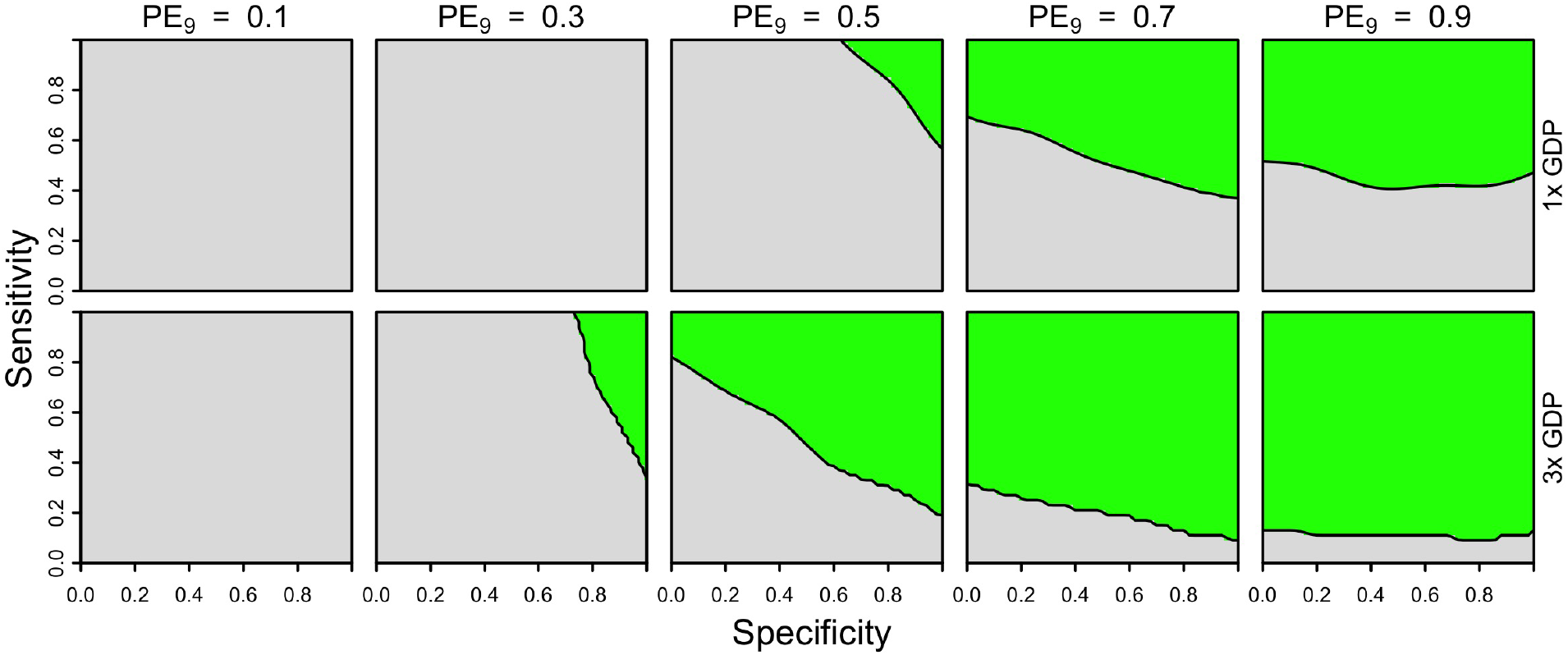
Cost-effectiveness of the intervention from a public payer perspective over a 30-year period, assuming one dose of vaccine (23 USD) and a fixed cost of serological screening (10 USD) under Brazil-like cost assumptions. Cost-effectiveness according to eqn. 5 is shown in green as a function of sensitivity (y-axis), specificity (x-axis), and PE_9_ value (columns). The value of cost_DALY_ is equal to per capita GDP (8,650 USD) in the top row and three times per capita GDP in the bottom row.

**Figure S19.**
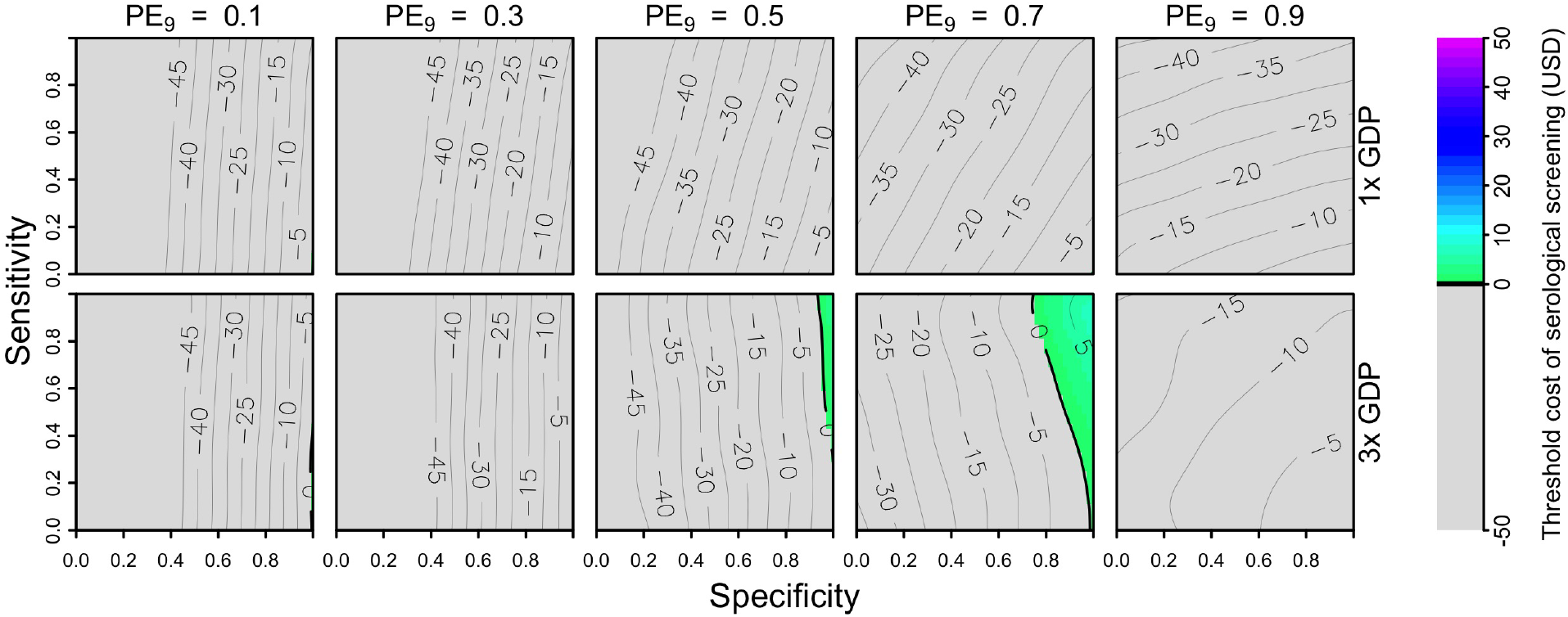
Threshold cost of serological screening from a public payer perspective over a 30-year period, assuming a vaccination cost of 69 USD and economic assumptions from the Philippines. Threshold costs are indicated by color as a function of sensitivity (y-axis), specificity (x-axis), and PE_9_ value (columns). The value of cost_DALY_ is equal to per capita GDP (8,650 USD) in the top row and three times per capita GDP in the bottom row.

**Figure S20.**
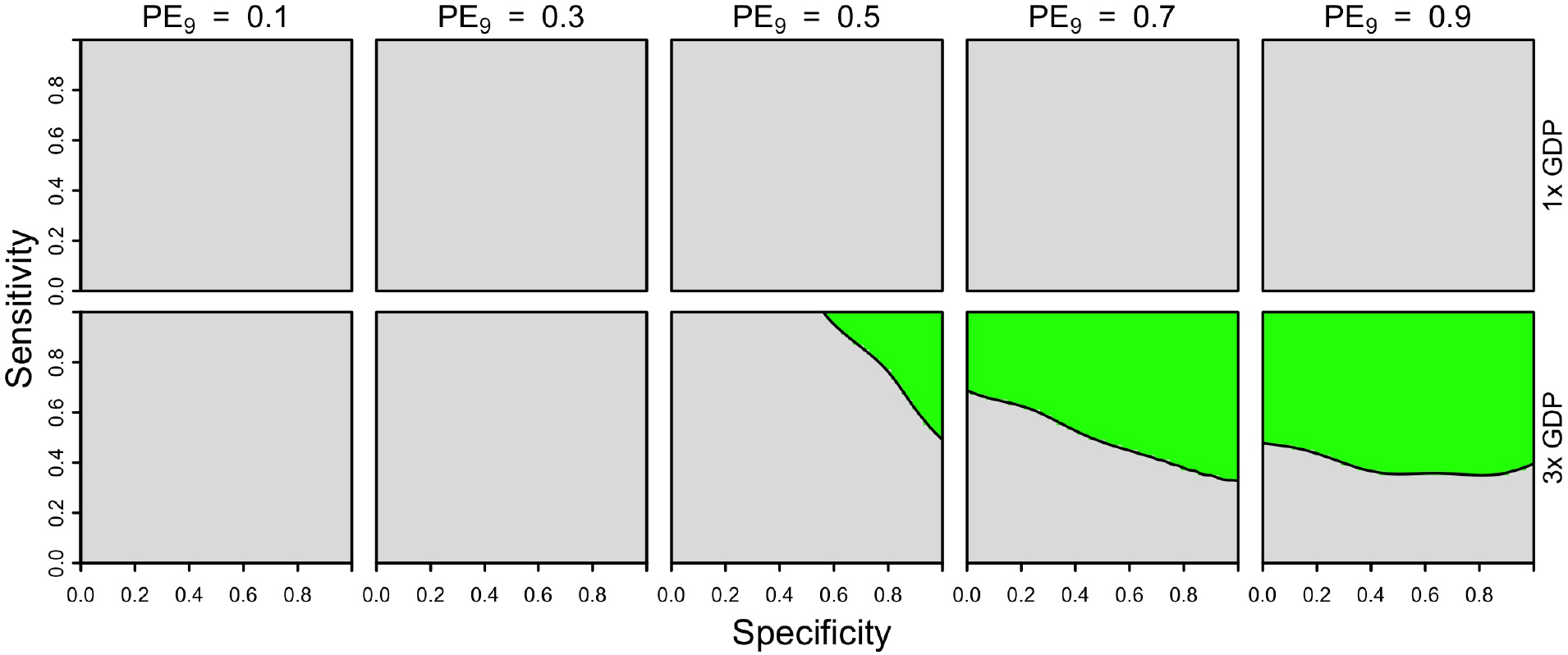
Cost-effectiveness of the intervention from a public payer perspective over a 30-year period, assuming one dose of vaccine (23 USD) and a fixed cost of serological screening (10 USD) under the Philippines-like cost assumptions. Cost-effectiveness according to eqn. 5 is shown in green as a function of sensitivity (y-axis), specificity (x-axis), and PE_9_ value (columns). The value of cost_DALY_ is equal to per capita GDP (8,650 USD) in the top row and three times per capita GDP in the bottom row.

**Figure S21.**
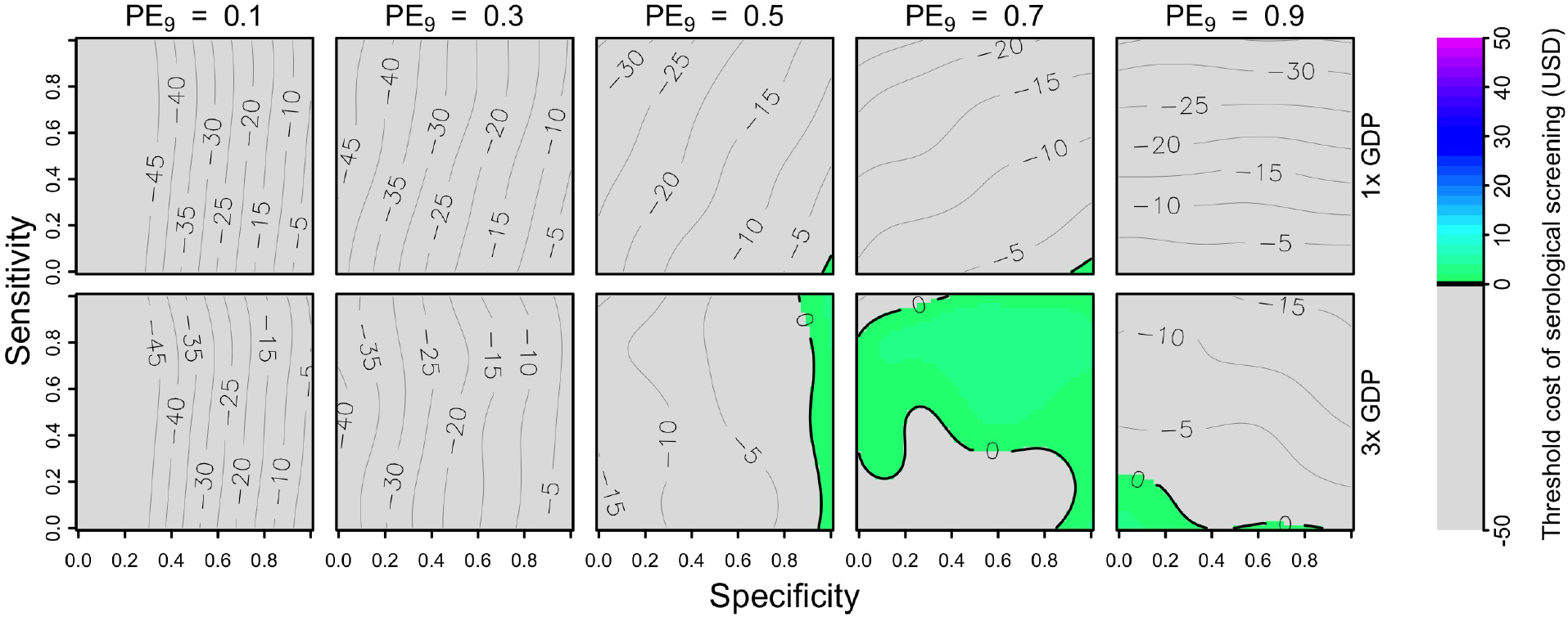
Threshold cost of serological screening from an individual perspective over a 30-year period, assuming a vaccination cost of 69 USD and economic assumptions from Brazil. Threshold costs are indicated by color as a function of sensitivity (y-axis), specificity (x-axis), and PE_9_ value (columns). The value of cost_DALY_ is equal to per capita GDP (8,650 USD) in the top row and three times per capita GDP in the bottom row.

**Figure S22.**
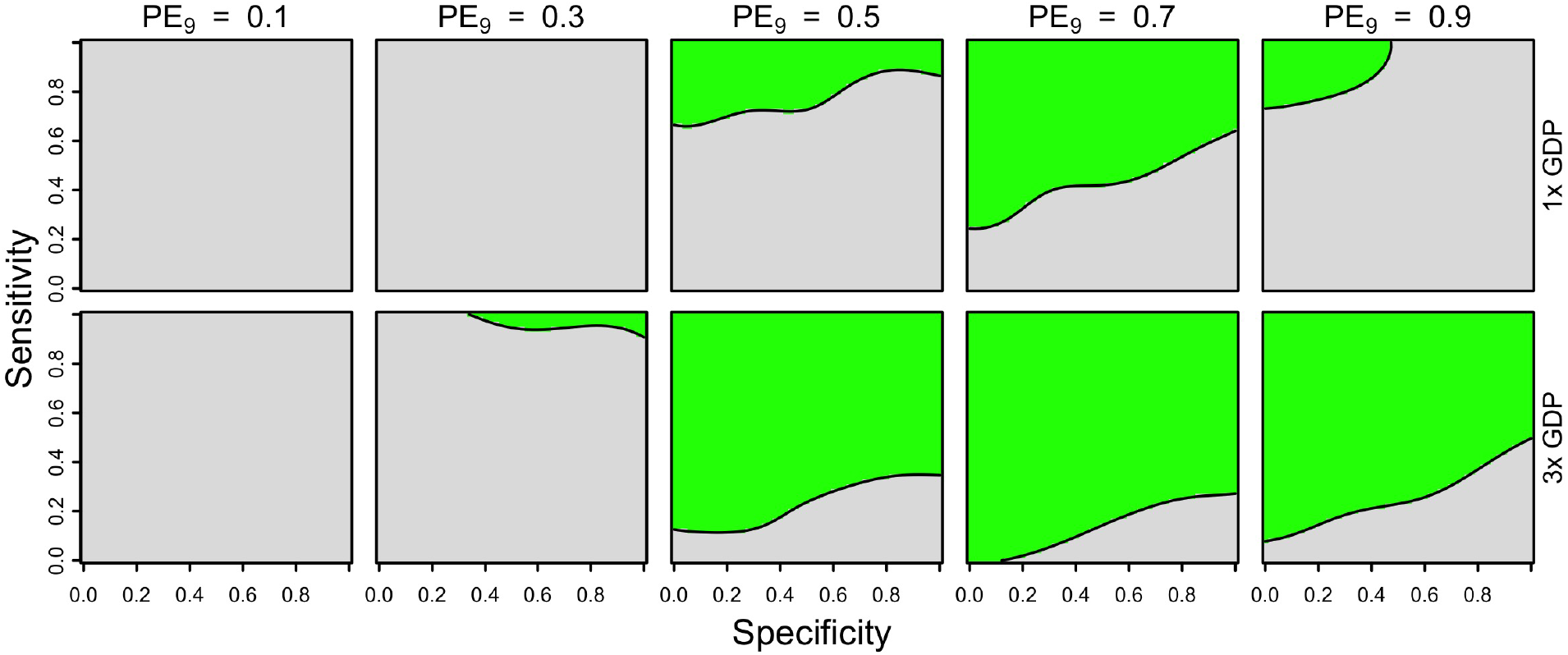
Cost-effectiveness of the intervention from an individual perspective over a 30-year period, assuming one dose of vaccine (23 USD) and a fixed cost of serological screening (10 USD) under Brazil-like cost assumptions. Cost-effectiveness according to eqn. 5 is shown in green as a function of sensitivity (y-axis), specificity (x-axis), and PE_9_ value (columns). The value of cost_DALY_ is equal to per capita GDP (8,650 USD) in the top row and three times per capita GDP in the bottom row.

**Figure S23.**
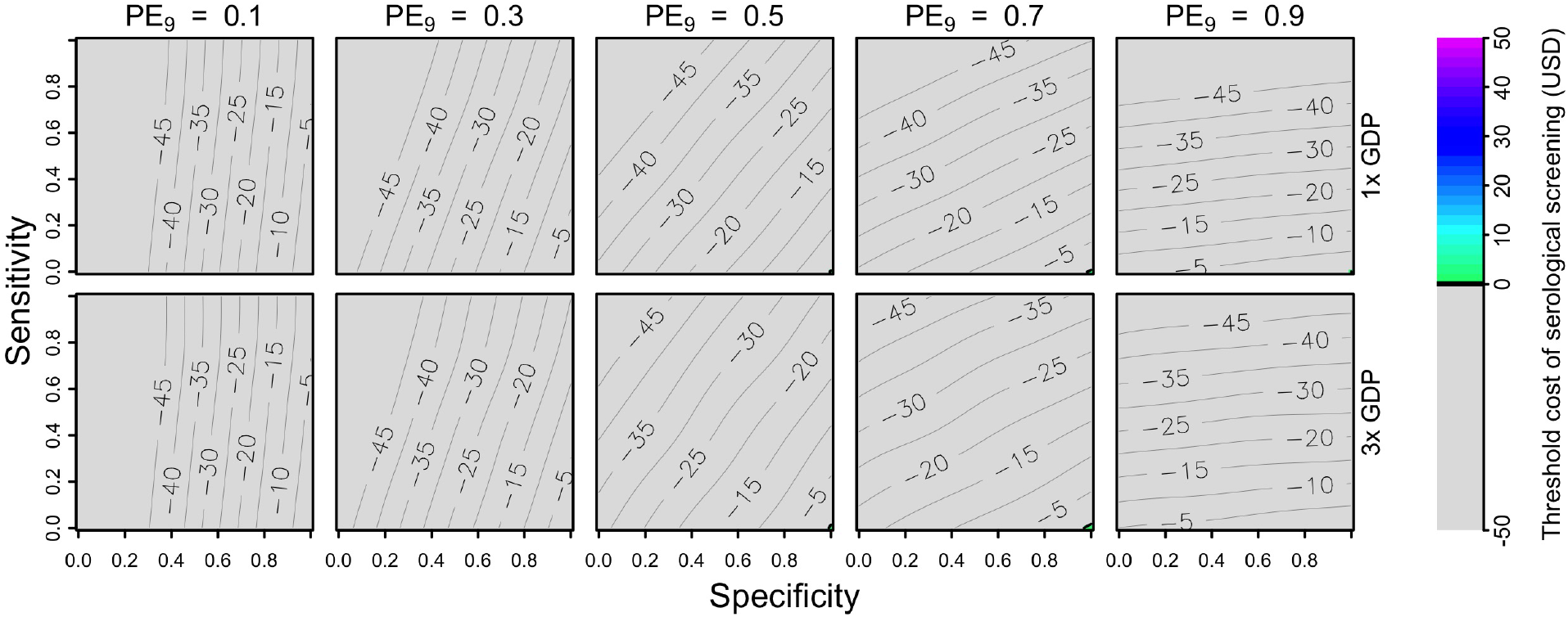
Threshold cost of serological screening from an individual perspective over a 30-year period, assuming a vaccination cost of 69 USD and economic assumptions from the Philippines. Threshold costs are indicated by color as a function of sensitivity (y-axis), specificity (x-axis), and PE_9_ value (columns). The value of cost_DALY_ is equal to per capita GDP (8,650 USD) in the top row and three times per capita GDP in the bottom row.

**Figure S24.**
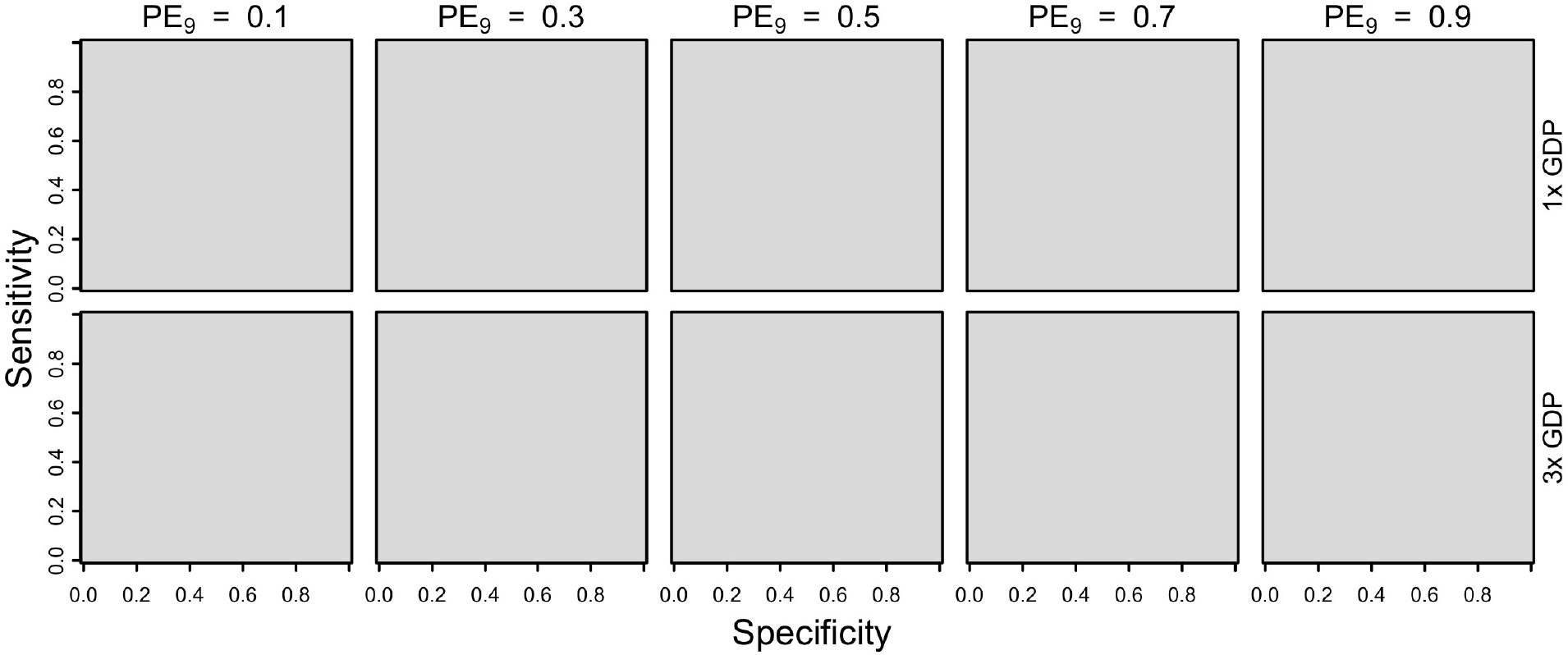
Cost-effectiveness of the intervention from an individual perspective over a 30-year period, assuming one dose of vaccine (23 USD) and a fixed cost of serological screening (10 USD) under the Philippines-like cost assumptions. Cost-effectiveness according to eqn. 5 is shown in green as a function of sensitivity (y-axis), specificity (x-axis), and PE_9_ value (columns). The value of cost_DALY_ is equal to per capita GDP (8,650 USD) in the top row and three times per capita GDP in the bottom row.

